# Analysis and culturing of the prototypic crAssphage reveals a phage-plasmid lifestyle

**DOI:** 10.1101/2024.03.20.585998

**Authors:** Danica T. Schmidtke, Angela S. Hickey, Ivan Liachko, Gavin Sherlock, Ami S. Bhatt

**Affiliations:** Department of Microbiology and Immunology, Stanford University, Stanford, CA, USA; Department of Genetics, Stanford University, Stanford, CA, USA; Phase Genomics, Seattle, WA, USA; Department of Medicine (Division of Hematology), Stanford University, Stanford, CA, USA

**Keywords:** crAssphage, microbiome, bacteriophage, plasmid, phage-plasmid, *Phocaeicola*, *Bacteroidota*, *Carjivirus communis*

## Abstract

The prototypic crAssphage (*Carjivirus communis*) is one of the most abundant, prevalent, and persistent gut bacteriophages, yet it remains uncultured and its lifestyle uncharacterized. For the last decade, crAssphage has escaped plaque-dependent culturing efforts, leading us to investigate alternative lifestyles that might explain its widespread success. Through genomic analyses and culturing, we find that crAssphage uses a phage-plasmid lifestyle to persist extrachromosomally. Plasmid-related genes are more highly expressed than those implicated in phage maintenance. Leveraging this finding, we use a plaque-free culturing approach to measure crAssphage replication in culture with *Phocaeicola vulgatus, Phocaeicola dorei,* and *Bacteroides stercoris*, revealing a broad host range. We demonstrate that crAssphage persists with its hosts in culture without causing major cell lysis events or integrating into host chromosomes. The ability to switch between phage and plasmid lifestyles within a wide range of hosts contributes to the prolific nature of crAssphage in the human gut microbiome.

## Introduction

Phages, viruses that infect prokaryotes, are among the most abundant genetic entities on earth, yet until the advent of affordable metagenomic sequencing, only a small number of human microbiome-related phages were known. Now, DNA viruses are estimated to outnumber bacteria in the human gut 2:1^1^ and certain phages, such as the prototypical crAssphage (*Carjivirus communis*), are thought to be among the most prevalent and abundant genetic entities associated with humans^2^. Despite these advances, human associated phages remain under-characterized and under-cultured despite their importance in shaping both microbial communities and human health^3–7^. Even after over a decade of study, *Carjivirus communis* has not been successfully propagated and has yet to be isolated in pure culture.Therefore, there is little understanding of its biology, lifestyle, and the implications of those on human health.

Phages in the gut likely impact human health both indirectly, by modulating the composition of the gut microbiota, and directly, through interactions with mammalian cells^8^. Phages can manipulate gut bacterial community composition by lysing their bacterial hosts leading to shifts in community composition that are often correlated with disease states^9–11^. Furthermore, phages can facilitate horizontal gene transfer between gut bacteria and can encode virulence factors such as toxins, which allow bacteria to more readily cause disease^8^. Finally, phages can directly interact with their human superhosts, as they have been shown in specific cases to transverse the gut epithelial barrier, enter the circulatory system, and inject their genomes into human cells where, sometimes, phage genomes can be transcribed^12,13^. The impacts that phages have on their hosts and super hosts are likely related to the phages’ lifestyles.

Classically, phages are categorized as either purely lytic or lysogenic. Purely lytic phages use host machinery to replicate and package their genomes into capsids followed by cell lysis to release phage progeny, killing their host in the process. By contrast, temperate phages can integrate into the host genome, only rarely transitioning into a lytic phase. In some environments, like the ocean, the majority of phages are thought to be lytic^14^. However, with advances in computational tools^15^ increasing the sensitivity of prophage detection, it is now recognized that the vast majority of phages that reside in the human gut are likely temperate^8^. Increased bacterial lysis (which can be caused by lytic phages) has been described to trigger disease states such as inflammation and increased gut permeability^8,13,16–19^. Taken together, the balance between lysogeny and lysis is likely important in maintaining human health ^8,20^. As more phages are discovered, it is becoming clear that many seemingly “temperate” gut phages do not encode classical marker genes of temperate lifestyles, such as integrases for integrating into bacterial genomes^21^. Therefore, alternatives to phage genome integration are likely common in the gut, allowing phages to persist for long periods as non-integrative lysogens ^21^.

One mechanism by which phages have evolved a non-integrative lysogenic state is through the acquisition of plasmid genes. Phage-plasmids (PPs) are large DNA elements (larger than phage or plasmid alone, >90Kb), that encode phage, plasmid, and accessory genes^22,23^. Rather than switching between lysis and integration, PPs switch between phage lysis and low-copy number plasmid replication modes^24^. A recent, large computational analysis showed that PPs are far more prevalent than previously appreciated, and that ∼7% of all sequenced plasmids and ∼5% of phages in the RefSeq database are likely PPs^22^. Even these numbers are likely gross underestimates of the true prevalence of PPs, because PP detection methods require searching for plasmid gene annotations in phage genomes and vice versa and gene annotations in phages are rather limited^25^. Due to their lifestyle, which includes (with some exceptions) recurrent productive infection, the lack of chromosomal integration, and a decreased propensity for plaque formation, few PPs are culturable^26–31^. Additionally, even cultured phages or plasmids might escape identification as PPs because one life cycle may dominate (phage or plasmid) thereby preventing experimental observation of both lifestyles in culture. Due to the challenges with identifying, culturing, and studying PPs, few PPs have been cultured or studied in depth, and we have only just begun to understand their prevalence and diversity.

*Carjivirus communis* is infamously difficult to culture, despite its high prevalence and abundance in the human gut. *C. communis’* 97Kb dsDNA genome was computationally discovered by metagenomic cross-assembly, and estimates for the global prevalence of *C. communis* are >70% with abundances reaching >90% of publicly available human gut viral-like-particle sequencing^2,32^. Despite rigorous attempts, *C. communis* is not known to grow on isolated bacteria, form plaques, integrate into bacterial chromosomes, or encode integration-related genes^2,33–37^. However, Guerin *et al.* demonstrated that the *Carjivirus* genus can replicate in a continuous stool culture suggesting that *C. communis* and its bacterial host are both abundant in stool and culturable^34^. Therefore, we hypothesize that, like PPs, *C. communis* might use an alternative lysogenic state that does not include integration into the bacterial host genome.

Here, we report the first culturing of *C. communis* on a single host, and classify it as a phage-plasmid. First, we identify genomic features that are consistent with a PP lifestyle. Next, we predict *Phocaeicola vulgatus* as a bacterial host for *C. communis* via proximity-ligation sequencing of *C. communis* containing stool. Through both proximity-ligation sequencing and long-read sequencing we are unable to observe integration of *C. communis* into bacterial genomes, but do observe the potential existence of both circular and linear forms of the *C. communis* genome. Given previously failed plaquing attempts, we investigate whether *C. communis* might predominantly exist as a plasmid. Analysis of publicly available metatranscriptomic data reveals that the *C. communis* plasmid genes are more highly expressed than phage genes in stool samples, suggesting that the plasmid lifestyle is preferred in the context of a healthy human gut. We then seek to culture *C. communis* to observe its growth dynamics. We develop a culturing approach to study *C. communis* and begin to characterize its phage-plasmid-like lifestyle, that enables targeted phage culturing and does not necessitate plaque formation. The culturing technique is as follows: 1) identification of phage containing stool, 2) bacterial host prediction via proximity ligation sequencing, 3) stool-based culture, 4) harvest phage filtrate from stool-based culture, and 5) liquid culture with the predicted host. We demonstrate that *C. communis* replicates not only in stool-based culture, but also in pure *P. vulgatus, Phocaeicola dorei* and *Bacteroides stercoris* cultures. Upon culturing *C. communis*, we observe growth dynamics that differ across hosts and a lack of plaque formation. Taken together, the high expression of plasmid genes, segments of time with stable phage:host ratios observed in culture, and lack of both integrative lysogeny and plaque formation suggest that *C. communis* is a phage-plasmid that predominantly exists in plasmid form.

## Results

### Genomic analysis of *Carjivirus communis*

Since few PPs are culturable, most knowledge surrounding PPs comes from the in-depth characterization of one PP, bacteriophage P1^22,24,27,28,38,39^. Given the absence of widely agreed-upon criteria for defining a PP, we used the key genomic and lifestyle features of P1 as a benchmark against which to evaluate *Carjivirus communis* as a candidate PP. When P1 is a phage, it has a linear, double-stranded DNA genome with terminally redundant sequences whose sequence similarity causes the genome to pseudo-circularize^24,26^. When P1 is a plasmid, the genome circularizes through a recombinase-mediated process, and it stably exists at one copy per bacterial chromosome via plasmid addiction with a toxin-antitoxin (TA) system and a partitioning system (ParA ATPase Walker)^24^. P1 also requires two different origins of replication, one for each lifestyle. P1 replicates as a circular plasmid from one origin (*oriR*) located within the replication initiation gene (repL) and, less often, induces its lytic cycle and replicates from a second origin (*oriC*) partially located within a second replication initiation gene (repA)^24^. Finally, P1 is prevalent in the gut, and due to its wide host range, and prolific transduction, it majorly contributes to horizontal gene transfer across diverse bacteria in the gut^38,40^. Here, we show that *C. communis*, like P1, is a PP with phage and plasmid gene functions, linear and circular genome forms, and two origins of replication.

### *Carjivirus communis* encodes both phage and plasmid genes

The *C. communis* genome has two distinct regions with opposing gene orientation and function^2^. The genes on the positive strand encode genome replication-related functions, while the reverse strand encodes phage structural and lytic-related genes (Fig. 1A). We were curious if this division of gene orientation and function permits two *C. communis* lifestyles; one “plasmid” and one “phage.” We hypothesized that *C. communis* might encode genes important for a plasmid lifestyle on the positive strand.

**Figure 1.**
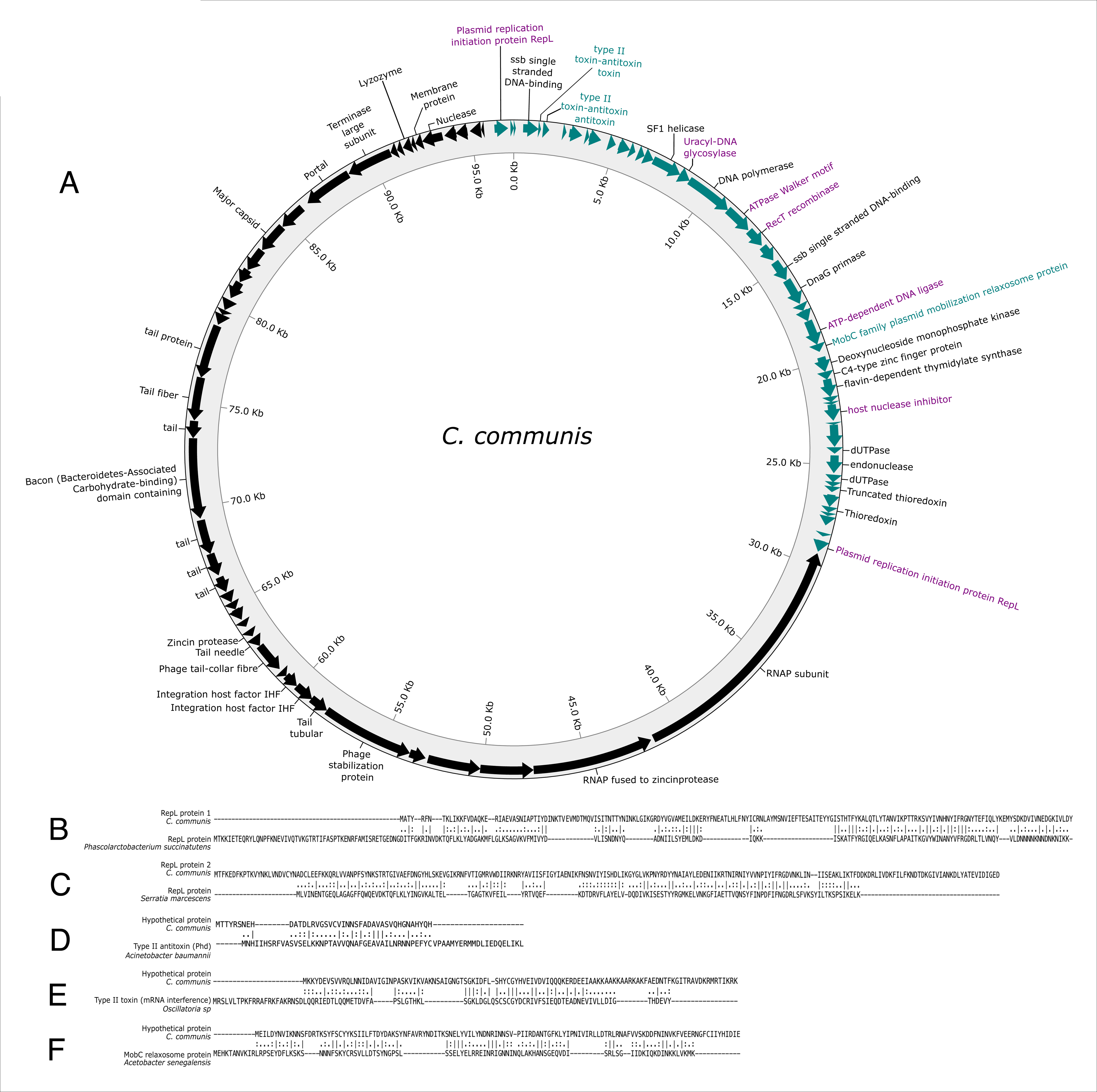
The *C. communis* genome encodes both phage and plasmid features. Visual representation of the *C. communis* reference genome with gene annotations and protein alignments of key genes A) The *C. communis* genome. Gene color denotes strand orientation (forward = green, reverse = black). Gene annotations are labeled; those in green originate from plasmids, those in purple are implicated in plasmid functions but did not show significant sequence similarity to any plasmid proteins. Text color of plasmid related genes denotes previously annotated genes (purple) versus gene annotated in this study (green). B/C) protein alignment of predicted RepLs in *C. communis* and their closest plasmid relatives. D) protein alignment of predicted antitoxin in *C. communis* and its closet plasmid relative. E) protein alignment of predicted toxin in C. communis and its closet plasmid relative. F) protein alignment of predicted MobC in *C. communis* and its closest plasmid relative.

While Dutilh *et al.* annotated two plasmid-originating genes in the original *C. communis* reference genome, these genes have not been further analyzed^2^. These two genes encode candidate plasmid replication initiation proteins (RepL) (Fig. 1B-C). One *repL* is in the first position of the positive-strand genes and the other is in the last position, and their protein sequences are only distantly related to one another (24.6% amino acid identity). The duplication of *repL* may allow for two origins of replication to regulate between phage and plasmid lifestyles. The presence of these genes also resulted in the identification of *C. communis* in a large computational analysis for PPs^22^. However, *C. communis* was the only *Crassvirales* found in the analysis and did not cluster with any other PPs and was not examined further.

Given that *Crassvirales* in general show low rates of integrative lysogeny and yet persist in the gut and laboratory cultures for long periods, we were curious if the PP lifestyle might be common across *Crassvirales.* We identified 23 *repL* genes in the genomes of 19 additional *Crassvirales* genomes (Supplementary Fig. 1). Five of these RepL proteins are encoded in four *Crassvirales* genomes (one copy in four of the genomes, and two copies in one of the genomes) with >95% nucleotide identity shared across >96% of the *C. communis* genome, while the rest of the RepLs are in more divergent *Crassvirales* genomes (Supplementary Fig. 1). Due to the diverse nature of *Crassvirales,* it is possible that *repL*-like genes are more widespread throughout *Crassvirales* genomes, but their sequences are too divergent to be easily identified.

While the presence of *repL* is strong evidence for a PP lifestyle, we nonetheless looked for other plasmid-like genes within the *C. communis* genome. To identify such genes we created a custom BLAST database of plasmid protein sequences and compared the *C. communis* protein sequences to the custom database^41^. We identified two additional genomic features that are present in other PPs; a potential TA system and a *mobC* gene (Fig. 1D-F). TA systems cause plasmid addiction and regulation of plasmid copy number, and the presence of a TA system in *C. communis* likely contributes to its persistence in the gut^42^. MobC allows plasmids to mobilize by hitchhiking through existing conjugative structures^43^. MobC is part of the relaxosome which binds the origin of transfer, melts the DNA, creates a single-stranded nick, and pulls the ssDNA into the new cell where it is then ligated into circular ssDNA and replicated to become dsDNA again^43–45^. In further support of this gene annotation, there is a previously annotated DNA ligase directly next to the predicted *mobC* gene (Fig. 1A). While we could not identify homology between this DNA ligase and other *mob* genes, it may function like MobA, another component of the relaxosome that has DNA ligase capabilities^43^.

Finally, two previously identified genes, a uracil-DNA glycosylase and a host nuclease inhibitor are likely implicated in anti-phage defense^46–51^. The presence of these genes also points towards a PP lifestyle since PPs tend to have larger genomes and encode more accessory functions such as a wide repertoire of counter mechanisms to bacterial-encoded phage defense and plasmid TA systems^22,39,52^.

### Bacterial host prediction for *Carjivirus communis* via proximity ligation sequencing

Given that PPs tend to exist as circular extrachromosomal elements, we first wanted to determine if *C. communis* is an integrative prophage or is maintained extrachromosomally. To search for *C. communis* integration into its bacterial host, we first had to determine a potential bacterial host. However, using strictly computational methods to predict the bacterial host(s) of phages within complex microbial communities is difficult, tends to have low accuracy, and does not always produce species-level predictions^53^. There are many computational tools for phage-host prediction, but most computational predictions have not been experimentally validated and tend to show inconsistencies across methods^53,54^. Additionally, most computational host prediction methods rely on assumptions that are not universally true, such as: GC content or methylation pattern matches between phage and their hosts, sequence matches to existing databases, phage integration into host genomes, and matched codon usage between phages and their hosts^53,54^. Previous computational host predictions largely agree that the host of *C. communis* is within the phylum *Bacteroidota* (Table 1), yet there is no obvious agreement as to which species may serve as a *C. communis* host. These inconsistencies in host predictions likely arise because most previous prediction methods were strictly computational and have low accuracy. It is also likely that the success of *C. communis* in the gut may be attributed to a wide host range and host predictions may never agree on a single bacterial host.

An alternative approach for phage-host identification, which has been shown to be quite accurate, is metagenomics proximity-ligation sequencing^55,56,56–61^. This technique captures phage-bacterial chromosome interactions in three-dimensional space, enabling the identification of phage genomes and bacterial genomes that are in very close physical proximity to one another (i.e., in the same cell; Fig. 2A). Specifically, the proximity of the bacterial and bacteriophage genomes enables crosslinking and ligation between the molecules, which then produces chimeric bacteria-phage DNA molecules that can be quantified as evidence of co-localization within a cell (Fig. 2A).

**Figure 2.**
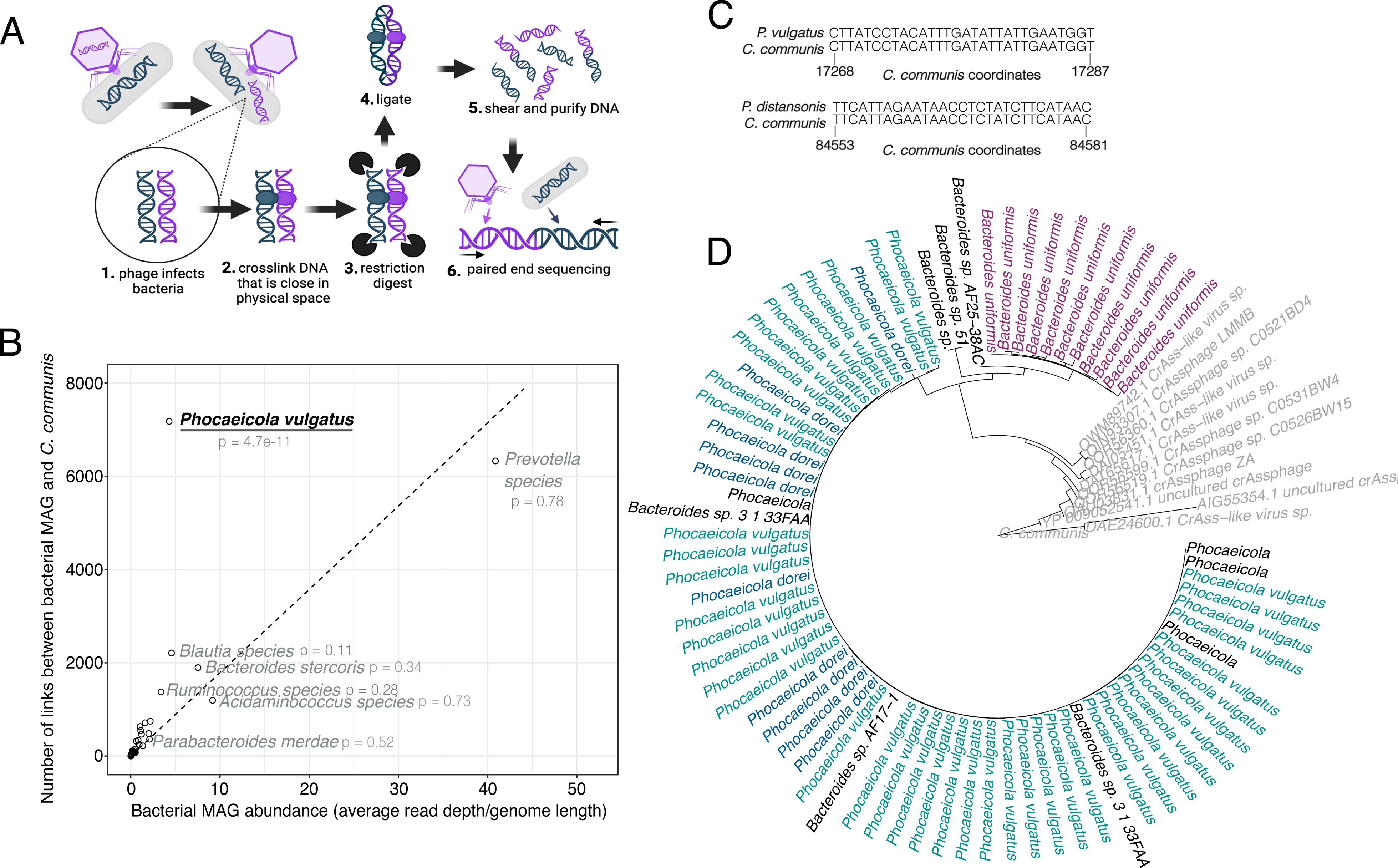
ProxiMeta Hi-C sequencing for bacterial host prediction of *C. communis*. A) Schematic of meta-Hi-C sequencing for host prediction of phages made in BioRender. B) plot of bacterial MAG abundance vs. chimeric *C. communis*-bacterial MAG reads (“links”) for host prediction of C. communis. Also, see supplementary figure 2. C) Clustered regularly interspaced short palindromic repeats (CRISPR) spacer analysis via PHISdetector for *C. communis* host prediction. D) phylogeny of blastp hits to the *Bacteroides*-associated carbohydrate-binding often N-terminal (BACON) domain containing protein from the *C. communis* genome for bacterial host prediction.

We thus applied ProxiMeta Hi-C for the identification of a candidate *C. communis* bacterial host. To identify a *C. communis* containing stool sample we analyzed previously published shotgun metagenomic sequencing data (for which we had matched stool) for the presence of *C. communis*^62^. For one sample, we found that 11.5% of the shotgun metagenomic reads mapped to the *C. communis* reference genome. We found that the *C. communis* genome was linked to *Phocaeicola vulgatus* more than expected by chance for a metagenome assembled genome (MAG) of its abundance (Fig. 2B). As a negative control, we also mapped the links between *C. communis* and two other MAGs present in the sample but not predicted as hosts (*Parabacteroides merdae* and *Bacteroides stercoris*). The observed links between *C. communis* and these two bacteria were much lower frequency than the internal links for each bacterial genome (Supplementary Fig. 2). By contrast, the frequency of links between *P. vulgatus* and *C. communis* was comparable to the internal *P. vulgatus* links (Supplementary Fig. 2), suggesting the links do not occur by chance. Importantly, our host prediction agrees with the only other experimental host prediction method that has been used for *C. communis* (single-cell microbiome sequencing)^63^ (Table 1).

### Bacterial host prediction for *Carjivirus communis* via CRISPR spacer analysis and gene homology

We additionally used CRISPR spacer analysis and phage-bacteria gene homology to predict a potential bacterial host for *C. communis*. PHISdetector^64^ revealed two perfect spacer matches in the *C. communis* reference genome, one to *P. vulgatus* and a second to *Parabacteroides distasonis* (Fig. 2C). Finally, the exchange of genetic material between phages and their bacterial hosts is common, and therefore searching for genes of bacterial origin within phage genomes can also point towards phage-host associations. One previously identified gene in *C. communis* is particularly suited for this task. A *Bacteroides*-Associated Carbohydrate-binding Often N-terminal (BACON) domain-containing protein was previously annotated in *C. communis* and is commonly encoded in *Bacteroidota* species^65^. Therefore, this sequence likely originated from a bacterium that *C. communis* previously infected. We determined that the *C. communis* reference BACON protein sequence was most closely related to proteins in other *Crassvirales*, followed by *P. vulgatus, P. dorei*, and *B. uniformis* (Fig. 2D). Synthesizing the results of these three orthogonal phage-host prediction methods, as well as the previous predictions from the literature, we predict that *P. vulgatus* is highly likely to serve as a host for *C. communis*, though it may not serve as the sole host.

### *Carjivirus communis* does not integrate into bacterial genomes

With a predominant bacterial host predicted, we sought to identify integration events of *C. communis* in its host. Therefore, we mapped the locations of the links between *C. communis* and *P. vulgatus* across the genomes (Fig. 3A) with the expectation that a random or even distribution across *P. vulgatus* suggests no integration, while a high density of links in one location would suggest *C. communis* integration at that location. We found an even distribution of links across the *P. vulgatus* genome suggesting a lack of integration of the phage into the bacterial genome (Supplementary Fig. 3).

**Figure 3.**
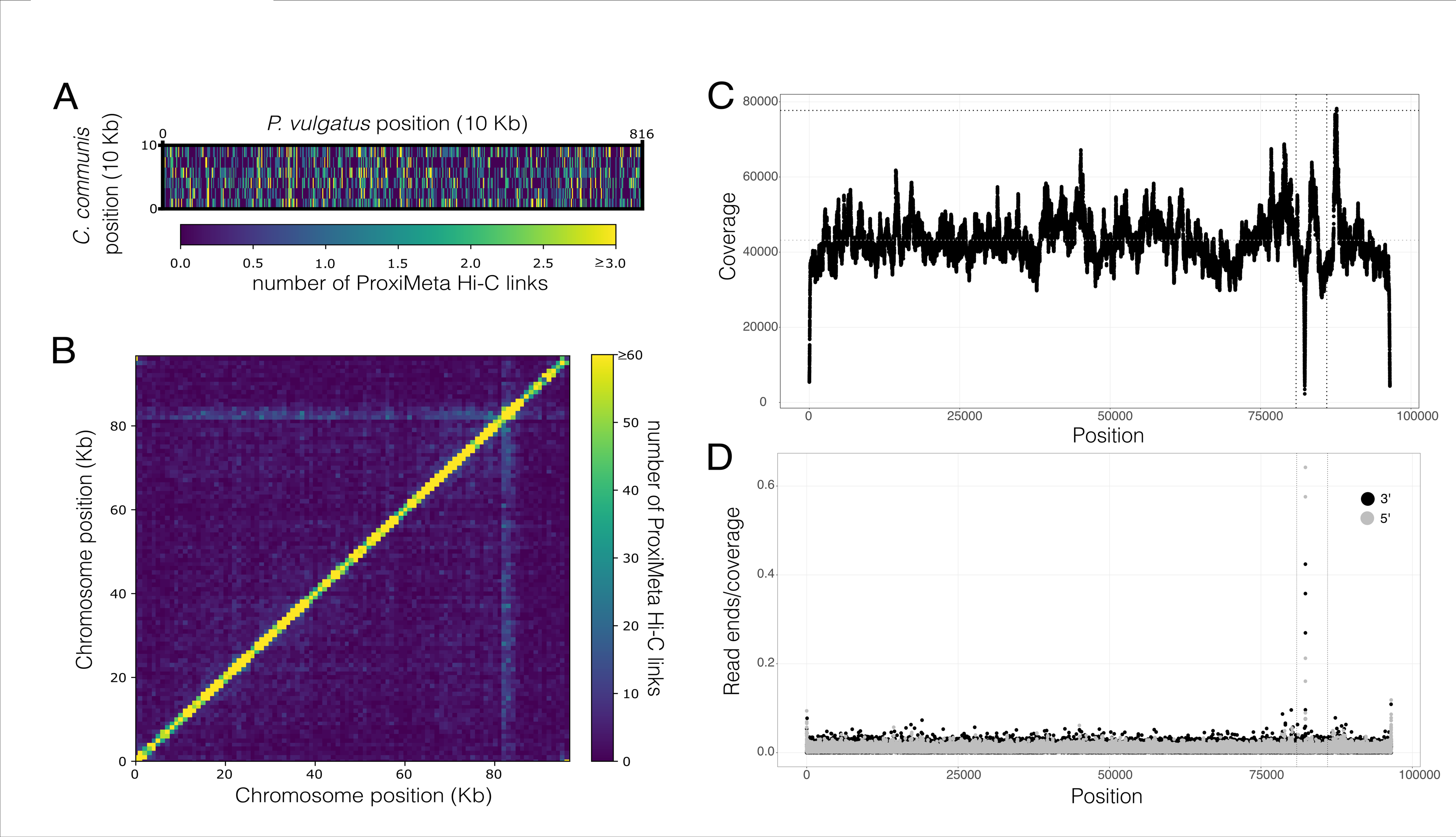
*C. communis* genome structure. A) ProxiMeta Hi-C linkage map of *C. communis* internal links, heat color represents number of ProxiMeta Hi-C links. Also, see supplementary figure 3. B) distribution of *C. communis*-*P. vulgatus* links across the *C. communis* genome (y-axis) and the *P. vulgatus* genome (x-axis), heat color represents the number of ProxiMeta Hi-C links. C) coverage (y-axis) of shotgun sequencing across the *C. communis* genome (x-axis). Gray dotted horizontal line represents average coverage, black dotted horizontal line is 1.8 times the average. Vertical dotted lines represent the boundaries of the global genome interactions in part B. D) Number of read ends (5’ in gray and 3’ in black) over coverage at that base (y-axis) across the *C. communis* genome.

Additionally, we analyzed long-read Oxford Nanopore sequencing of a *C. communis* positive stool for signatures of prophage integration. We identified reads that mapped to the *C. communis* reference genome and extracted soft-clipped read regions from those reads that had them. Extracted soft-clipped regions were then aligned back to the *C. communis* genome assembled from the sequenced sample: 99.9% (of 92,394 total) of the soft-clipped regions aligned directly back to *C. communis*, and the soft-clipped regions of only 18 reads remained unmapped. Of these, 12 had direct BLAST hits to or aligned to assembled contigs with hits to *Crassvirales* genomes, and the remaining 6 had no similarity to sequences in the database, and did not map to any assembled contigs from the sequencing; it is therefore unlikely that *C. communis* is integrated into any bacterial genomes in this dataset. These findings are consistent with prior efforts that also failed to find any evidence of *C. communis* integrative-prophages or lysogenic genes and observed overall very low rates of *Crassvirales* lysogeny compared to other gut phages^2,36^.

### *Carjivirus communis* exists as both a circular and a linear extrachromosomal element

Given that *C. communis* likely does not exist as an integrated lysogen, we sought to determine if it might be maintained extrachromosomally as a non-integrative lysogen. We hypothesized that *C. communis* predominantly exists as an extrachromosomal, circular plasmid, and less often as a linear phage, similar to P1^24^. First, we examined the ProxiMeta Hi-C data for interactions within the *C. communis* genome (Fig. 3B). The *C. communis* interaction map suggests predominantly standard, local interactions of a circular genome^66^. However, it also demonstrates global interactions between one segment of the genome and all of the other loci. A possible explanation for this observation is that if a linear version of the genome exists, the ends are more likely to partake in global interactions with the rest of the genome due to the increased bending of linear DNA allowing these interactions to take place more easily in three-dimensional space^67^.

Using previously established methods for phage genome terminus identification, we aligned our short read sequencing reads to our *C. communis* genome assembly and looked for regions of higher coverage (∼2x) which might suggest direct terminal repeats (DTRs) (Fig. 3C)^68^. We identified a region of high coverage in the same region of the genome that partakes in global genome interactions. Finally, we searched for read ends, and also identified a higher abundance of read ends at the global interaction region suggesting that the genome exists linearly (Fig. 3D). These data support the prior speculation that the genome may exist in two forms; linear with DTRs, and circular^69^. Additionally, *C. communis* encodes a RecT recombinase that might mediate the transition from a linear to a circular genome (Fig. 1A).

### *Carjivirus communis* encodes two origins of replication

The model phage plasmid P1 encodes two bidirectional origins of replication, one for plasmid replication initiation and one for lytic phage replication initiation, therefore we asked how *C. communis* might initiate replication^24,70^. Since *C. communis* encodes two RepLs we hypothesized that one RepL acts as an initiation factor for phage replication and the other other for plasmid replication. We used OriFinder^71^ to computationally predict an origin of replication in *C. communis*. OriFinder predicted a potential origin directly upstream of the *repL* at the beginning of the forward strand (Fig. 4A). RepL binds to an origin of replication, which in P1 is within the *repL* gene, to initiate genome replication. Therefore, the proximity of the *repL* gene and the predicted origin is supportive of the predicted origin location^24^. Additionally, OriFinder predicted a second potential origin ∼2,600 bp downstream of the other origin and directly downstream of the TA system. The TA system in P1 regulates plasmid copy number via competitive binding inhibition, and overproduction of the antitoxin allows it to bind to the origin in place of the replication initiation protein thereby inhibiting replication in a feedback loop to keep plasmid copy number under control.

**Figure 4.**
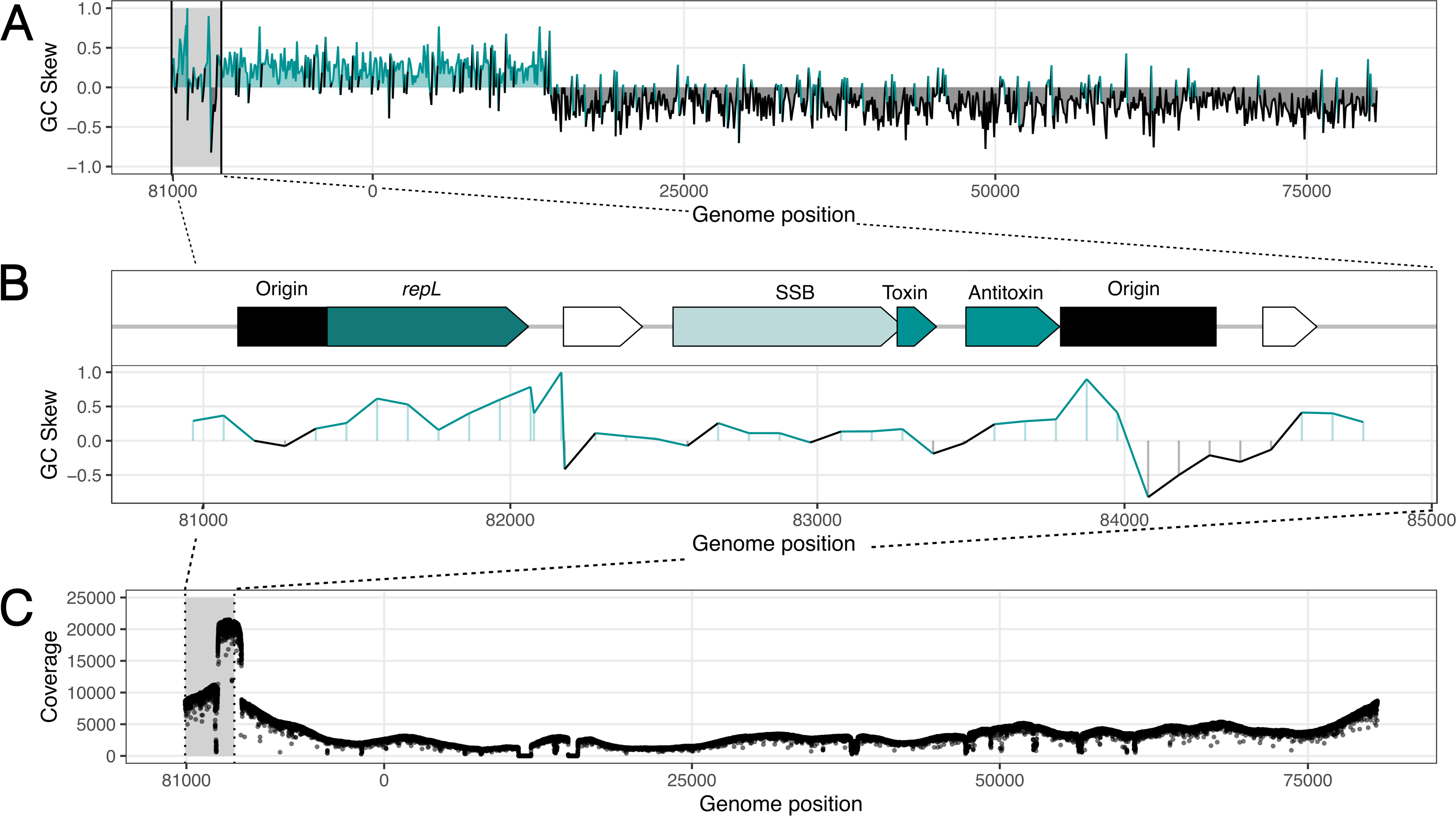
Two putative origins of replication in *C. communis*. A) GC skew plot of the entire *C. communis* genome. B) Genetic context of the two predicted origins of replication (green), aligned to the GC skew in that region of the genome. C) Coverage plot of Oxford Nanopore reads across the *C. communis* genome

The previously reported GC skew of the genome also supports the presence of both of these origins^72^. There is a stark shift in GC skew between the forward and reverse strands for the *C. communis* reference due to the perfect gene orientation coordination between the two strands. However, similar to the P1 genome, the GC skew between the two potential origins is close to zero^24^ (Fig. 4B).

Finally, we sought experimental evidence to explore whether the origins of replication might exist at the predicted region. Because replication firing at the origin is typically associated with a higher total copy number of DNA at the origin compared to the terminus or other parts of the genome, sequencing coverage is a potential readout for the origin location. The expectation is that coverage is the highest at the origin of replication and declines with distance from the origin. Therefore, we analyzed the coverage of the *C. communis* genome in long-read Oxford Nanopore sequencing, which shows higher coverage near the predicted origins of replication which declines further away from these loci (Fig. 4C). We conclude that *C. communis* likely uses two origins of replication similar to P1. P1 replicates bidirectionally from oriL early in its lytic phase, later transitioning to rolling circle replication thereby producing long linear concatemers of the genome. These concatemers are processed and packaged into phage heads as linear genomes with DTRs^24,70^. Once the phage infects a bacterial cell the P1 genome is circularized via recombination between DTRs and replicates as a plasmid from oriR until it again undergoes lytic replication^24,70^. Given the increasing evidence supporting the existence of both a phage and plasmid lifestyle for *C. communis*, and the lack of plaque formation or major lysis events, we investigated whether *C. communis* exists predominantly in a plasmid lifestyle.

### Plasmid genes are more highly expressed than phage genes in *Carjivirus communis*

To determine whether the phage or the plasmid genomic ‘program’ dominates when *C. communis* is in the human gut, we analyzed publicly available, paired stool metatranscriptomic and metagenomic data^73^. We identified 111 metatranscriptomic samples with >1x coverage of the *C. communis* genome. We calculated the mean expression for each individual gene in the *C. communis* genome across the 111 samples and found that genes on the forward (plasmid) strand had higher expression on average across all samples than those on the reverse (phage) strand (Fig. 5A). Next, we calculated the ratio of average positive stranded gene expression to negative stranded gene expression in each of the individual 111 samples. We found that the positive:negative ratio was >1 in 95/111 (85.6%) analyzed samples (Fig. 5B) suggesting that plasmid genes are more highly expressed in the majority of samples evaluated.

**Figure 5.**
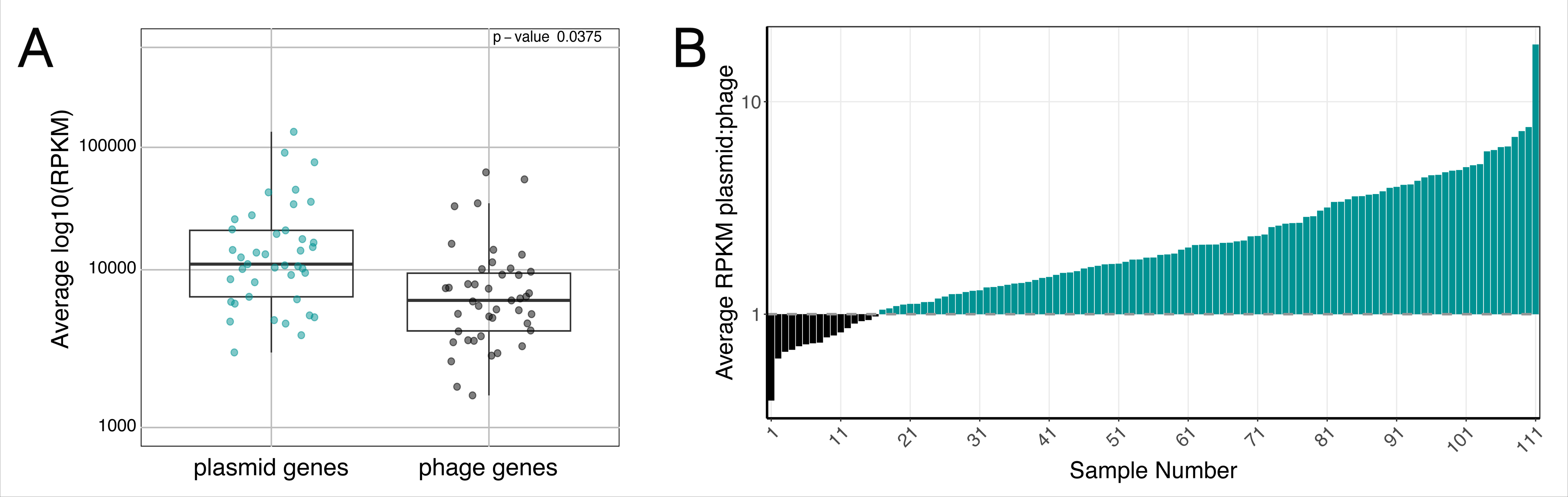
Expression of *C. communis* genes in metatranscriptomics. A) Average expression of each gene in the *C. communis genome* across 111 metatranscriptomics samples. Plasmid stranded genes in green, phage stranded gene in black. B) Ratio of average plasmid gene expression:average phage gene expression in each of the 111 samples. Also, see supplementary figure 4.

Finally, we examined the expression of *C. communis* genes across many samples. We looked for genes expressed in at least 105 of the 111 samples (∼95%). We found that genes encoding single-stranded binding proteins were the most often expressed (Single stranded binding protein 1 = 110/111, Single stranded binding protein 2 = 109/111) and were expressed at the highest average levels. The other genes expressed in the largest number of samples were a recombinase, RecT (110/111), SF1 helicase (109/111), ATPase walker motif (105/111), RNAP (105/111), and major capsid (105/111) (Supplementary Fig. 4, Supplementary Table 1). Interestingly, many of these genes are likely important in plasmid replication and concatemer resolution^24^. However, the major capsid is also expressed in many samples suggesting the production of phage particles. In addition to the major capsid, despite the lack of plaque formation by *C. communis,* we do observe expression of cell lysis and other structural genes suggesting that *C. communis* not only forms viral particles but also lyses out of bacterial cells under at least some circumstances.

### Targeted, plaque-free culturing of *C. communis*

A) *C. communis* has been heavily studied via computational analyses; however, further investigation into its biology requires culturing it in the laboratory. While we find supporting computational evidence that *C. communis* follows a PP lifestyle, we wanted to see if we could culture *C. communis* and observe PP-like growth dynamics. Therefore, we developed a plaque-independent culturing method that is well-suited for PP culturing. Additionally, phage culturing techniques typically entail isolation of many phages on a single bacterial host, rather than targeted isolation of a specific phage, and our culturing method allows targeted culturing of *C. communis.* The culturing technique includes: 1) identification of *C. communis* positive stool, 2) bacterial host prediction using ProxiMeta Hi-C, 3) stool-based culture, 4) harvesting phage filtrate from stool-based culture, and 5) liquid culture of *C. communis* with the predicted host.

### Identification of a *Carjivirus communis* strain in a stool sample

To culture the phage, we first needed to identify a reservoir from which we could replicate it. Given that the *Carjivirus* genus can replicate in continuous stool culture, we decided to use stool as a culturing source^34^. Mapping of metagenomic sequencing reads to the *C. communis* reference genome suggests a high abundance of a *C. communis* relative in the human stool sample on which we performed ProxiMeta Hi-C. To determine how closely the phage in this sample was related to the *C. communis* cross-assembled reference, we generated a complete genome using standard shotgun as well as ProxiMeta-Hi-C sequencing data for this stool sample. We compared our assembly to the *C. communis* reference genome and found that they are 95.5% identical (average nucleotide identity) over their entire lengths (Fig. 6), confirming that they are the same species^74,75^.

**Figure 6.**
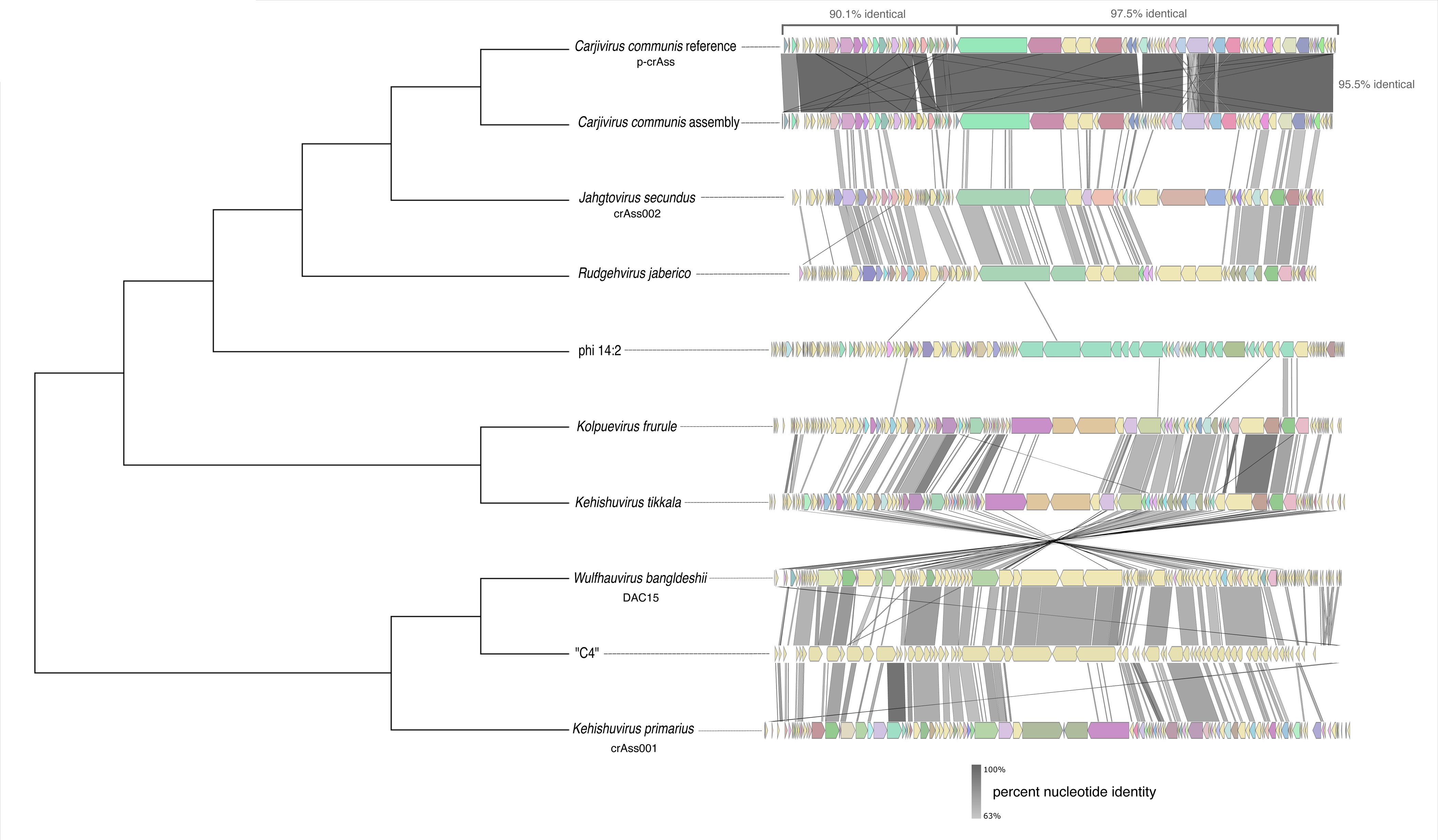
Relatedness of cultured *Crassvirales*. Whole genome alignment-based tree showing single representatives of cultured *Crassvirales* species, the *C. communis* reference genome, and the *C. communis* genome assembly the stool sample of interest is shown on the left. Synteny plots are shown on the right. The gray color scale connecting genomes in the synteny plots represents percent nucleotide identity between the genomes. Genes with identical annotations are colored the same. Hypothetical genes are yellow.

When comparing two distinct regions in the *C. communis* genome, separated by opposing gene orientation and coordinated function, we observed that the forward-stranded genes implicated in genome replication were overall less conserved (90.1% nucleotide identity) than the reverse strand implicated in phage particle production and cell lysis (97.5% nucleotide identity) between the *C. communis* reference genome and our assembly (Fig. 6). This observation is consistent with findings that phage genes are more highly conserved than their plasmid counterparts in PP genomes^22^. This suggests that while they may predominantly exist as plasmids, phage genes are likely not in the process of pseudogenizing.

Finally, we compared our assembled *C. communis* genome to the genomes of all previously cultured *Crassvirales* via whole genome alignment, construction of a phylogenetic tree, and generation of synteny plots to visualize similarities in gene organization (Fig. 6). Together, these data show that our assembly is a closer relative to *C. communis* than other cultured *Crassvirales* thus far. Having identified a stool sample with a strain of *C. communis*, and a potential bacterial host, *P. vulgatus,* we wanted to measure *C. communis* replication in relation to *P. vulgatus* in stool-based culture and observe phage-host growth dynamics

### Plaque-independent measurement of phage replication in stool-based culture

We sought to obtain a high-titer stock of the phage to then harvest and test for replication on the predicted bacterial host. Since it was previously reported that other phages in the *Carjivirus* genus can replicate in continuous stool culture, we first attempted to replicate *C*. *communis* to high titers in stool-based culture as a reservoir for further culturing^33,34^. However, since continuous culture systems are highly resource-intensive, we employed a non-continuous, batch culture system. We anaerobically cultured the *C. communis* positive stool sample identified above, alongside a second *C. communis* negative stool sample for which 0 shotgun sequencing reads mapped to the *C. communis* genome. Previous work found that the presence of kanamycin and vancomycin enriches for *Bacteroidota* and enhanced replication of the *Carjivirus* genus^34^. Therefore, we diluted our two stool samples into TSB media containing vancomycin and kanamycin and sampled the cultures every 4 hours across a 44-hour culture.

At each time point, we measured the OD600 of the culture (Fig. 7A) and observed robust growth of both stool samples. Additionally, since we could not recover *C. communis* plaques, we used qPCR to determine the copies of both the *C. communis* genome over time in the culture (Fig. 7B). Since the stool culture was a complex mixed community of bacteria, it was not possible to determine the abundance of the predicted bacterial host of *C. communis* (*P. vulgatus*) by colony-forming units (CFU), and we thus also measured genome copies of *P. vulgatus* via qPCR over time in the culture (Fig. 7B). We observed that the *C. communis:P. vulgatus* ratio settles around ∼1:1 from 24 to 32 hours followed by a ∼1.5 log increase in *C. communis* copies and an increasing *C. communis:P. vulgatus* ratio. A 1:1 ratio suggests either integration into a host genome, or extrachromosomal maintenance at a consistent copy number (n=1) per cell, like a plasmid. Since we do not observe any evidence of *C. communis* forming integrative lysogens, it likely exists as a plasmid.

**Figure 7.**
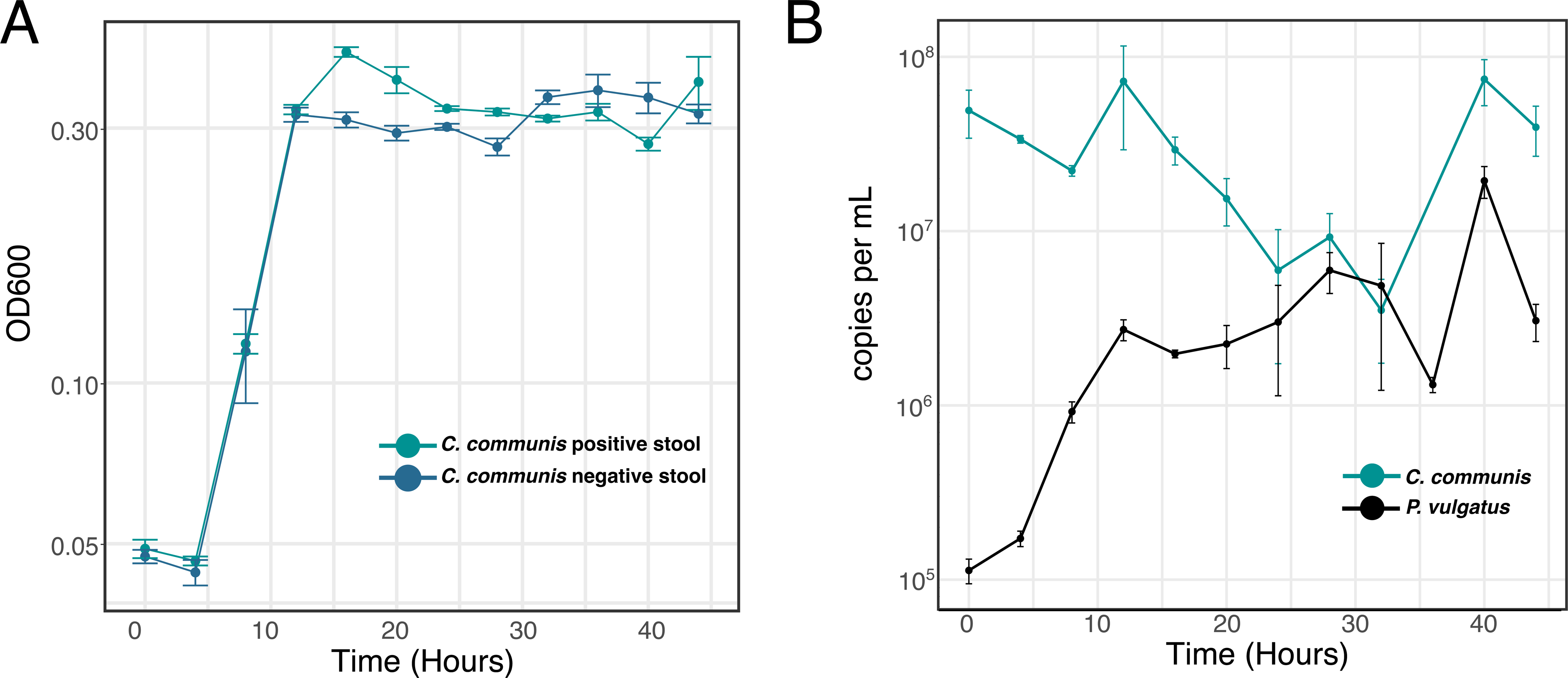
*C. communis* replication in stool-based culture. A) OD600 of a stool sample containing *C. communis* and a negative control stool sample that does not contain *C. communis*. B) copies of *C. communis* (green) and its predicted host, *P. vulgatus* (black), overtime in stool-based culture of *C. communis* positive stool as measured by qPCR. Both *P. vulgatus* and *C. communis* were undetectable by qPCR in stool-based culture of *C. communis* negative stool. Also, see supplementary figures 6-9.

### Experimental validation of *P. vulgatus* as a host for *C. communis*

Having demonstrated that *C. communis* can replicate in stool-based culture, we next sought to determine whether *C. communis* could 1) replicate on its predicted host, *P. vulgatus*, and 2) if it has similar growth dynamics in *P. vulgatus* pure culture as in stool culture. We first tested the ability of *C. communis* harvested in the phage filtrate from a fecal culture to plaque on the type strain *P. vulgatus* 8483 ATCC, as well as a strain that we isolated from stool which is closely related to *P. vulgatus*, *P. dorei.* Consistent with previously published work, we were unable to obtain *C. communis* plaques by standard methods^2,33^. Given the inducible nature of lysogenic phages, we also attempted to induce plaque formation through carbadox, mitomycin C, UV, and heat treatments, which similarly yielded no plaques on either bacterial species^31,38,39^. The lack of plaque formation, induction, and the observed ∼1:1 phage:host ratio in stool culture supports our model that *C. communis* might be largely maintained extrachromosomally at a stable copy number like a plasmid. Therefore, we turned to a liquid culturing approach.

We grew either *P. vulgatus* or *P. dorei* to mid-logarithmic growth and applied phage filtrate from the stool culture at roughly a multiplicity of infection (MOI) of 0.1 for *C. communis*. We sampled the culture every 4 hours over a 44-hour culture measuring the OD600 of the culture, CFU of the bacteria, and the copies of *C. communis* via qPCR each time point (Supplementary Fig. 6-9, Fig. 8A-B). In a liquid culture of a lytic phage, our expectation was that as the phage replicated, cells would lyse, and the OD600 of the culture would decline. However, we did not observe any difference between the OD600 of the bacterial cultures with and without phage filtrate added, despite the fact that *C. communis* replicated to high titers. We observed that *C. communis* copies increased by ∼1.5 logs when cultured with *P. vulgatus* and ∼3 logs with *P. dorei*.

**Figure 8.**
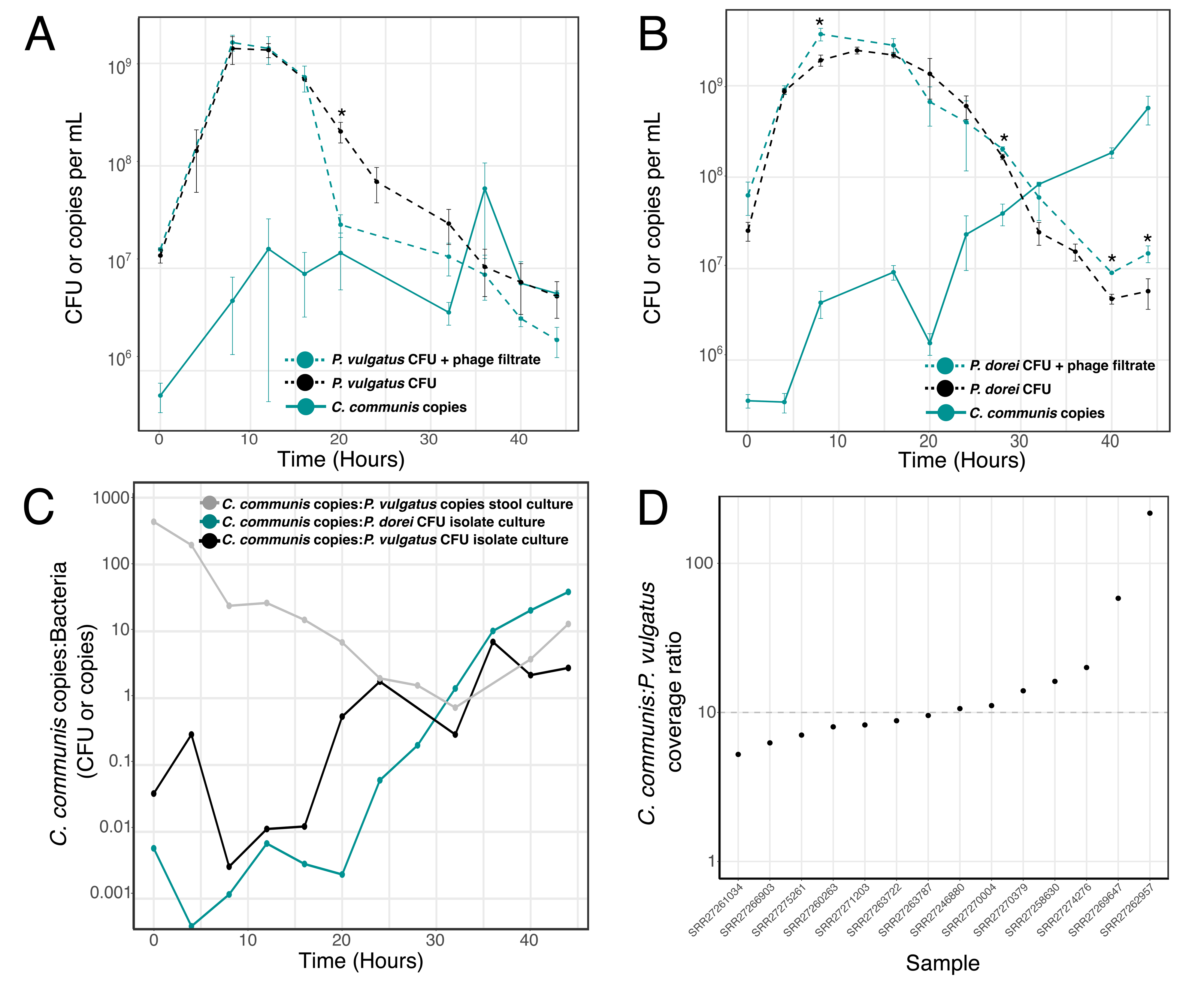
*C. communis* replication in isolate bacterial cultures. A) Copies of *C. communis* per mL of culture measured via qPCR (green solid line), CFU per mL of culture of *P. vulgatus* with phage filtrate (green dotted line) and without phage filtrate (black dotted line) * denotes that CFU per mL with and without phage filtrate added are statistically significantly different (p-value <0.05). (in the -phage filtrate condition copies of *C. communis* are undetected) B) Copies of *C. communis* per mL of culture measured via qPCR (green solid line), CFU per mL of culture of *P. dorei* with phage filtrate (green dotted line) and without phage filtrate (black dotted line) * denotes that CFU per mL with and without phage filtrate added are statistically significantly different (p-value <0.05). (in the -phage filtrate condition copies of *C. communis* are undetected) C) Ratio of *C. communis:P. vulgatus* in stool culture (gray), and *P. vulgatus* culture (black). Ratio of *C. communis:P. dorei* in *P. dorei* culture (green). D) *C. communis:P. vulgatus* ratio of genome length corrected coverage in publicly available single-cell microbiome sequencing. Also, see supplementary figure 5, 6, and 8.

As a negative control, we also added the same amount of phage filtrate to media only, to ensure that there was no carryover of the bacterial host from stool culture; we observed no bacterial growth and no significant increase in *C. communis* copies (Supplementary Fig. 6-9). Additionally, we tested for *C. communis* replication on two bacterial species present in the stool sample, but not predicted as hosts by ProxiMeta-Hi-C, *Parabacteroides merdae,* and *Bacteroides stercoris*. We did not observe significant *C. communis* replication in culture with *P. merdae,* but we did observe replication with *B. stercoris* (Supplementary Fig. 6-9). This suggests that *C. communis* has a wider host range than previously cultured *Crassvirales*, but that it does not ubiquitously infect all bacterial species in the stool sample that we cultured it from.

The ability of *C. communis* originating from phage filtrate to replicate on an isolated bacterium suggests that *C. communis* phage particles are produced in fecal culture, despite the lack of plaques and the observed ∼1:1 phage:host ratio in stool culture. This observation might point towards the ability of *C. communis* to exist in two separate lifestyles. Therefore, we were curious what the ratios of phage:host were in each culture and if we could observe ratios that suggest two different lifestyles. We observed different growth dynamics of *C. communis* in the fecal, *P. vulgatus*, and *P. dorei* cultures (Fig. 8C). Consistent with our findings in stool-based culture, we observed that when *C. communis* is grown on *P. vulgatus* the ratio of *C. communis* to *P. vulgatus* is roughly 1:1 from 24 to 32 hours of culture and remains between a 1:1 and 1:10 ratio for the remainder of the culture. In the *P. dorei* culture, we observed a steady increase in the *C. communis:P. dorei* ratio throughout the entirety of the culture. The difference in *C. communis* growth between with *P. vulgatus* vs. *P. dorei* suggests that the copy number of *C. communis* is not as well regulated in *P. dorei* as *P. vulgatus.* This might be attributable to the predicted TA system. If the TA system is adapted for growth on *P. vulgatus* but not adapted to *P. dorei*, *C. communis* might be able to reach higher copy numbers in individual *P. dorei* cells. The TA system may also drive *C. communis* to infect a high percentage of the total bacterial population through its plasmid addiction mechanism. However, in analyzing phage:host ratios we could not make conclusions about the percent of the bacteria that are infected at any given time point and therefore could not definitively conclude that there are two distinct lifestyles taking place.

To further explore the percentage of infected bacterial cells at any given time, and how many copies of *C. communis* might be present per cell, we analyzed publicly available single-cell microbiome sequencing data^63^. 14 single-amplified genomes (SAGs) of *P. vulgatus* were previously identified as *C. communis* positive. Thus, in these 14 samples, we determined the difference in coverage across the *C. communis* genome versus each *P. vulgatus* SAG as a readout for the copies of phage per bacterial cell (Fig. 8D). We found that there was a median ∼10:1 coverage of *Carjivirus communis:P. vulgatus* in the 14 samples, although 12 of these data range from ∼5-20 copies per cell, and there are two outliers. This is the same ratio observed at the final two time points in the *P. vulgatus* culture suggesting the possibility that 100% of cells are infected at this time point with roughly 10 copies of *C. communis* each. By contrast, two of the single cells had substantially higher copies of *C. communis*, >200 and ∼60, suggesting that *C. communis* may be in a phage-like lifestyle in these cells and that the burst size of *C. communis* may be ∼60-200 copies of phage per bacterial lysis event. Consistent with our single-cell analysis for *C. communis,* the burst size estimates for other cultured *Crassvirales* are ∼20-160 viral particles per cell^76^. One limitation to consider when interpreting these data is that often SAGs are generated through rolling circle amplification, which has been described to more efficiently amplify small circular DNA compared to larger circles or linear DNA^77^. Combining the evidence above, we conclude that *C. communis* may be capable of existing in two different lifestyles, one in which it maintains stable copy number as a plasmid and one in which it produces phage particles and lyses out of cells.

## Discussion

The 2014 discovery of *C. communis* led to the classification of a large, diverse, ubiquitous, and persistent order of phages, the *Crassvirales*^78^. *Crassvirales* are among the most heavily studied gut phages due to their incredible abundance and persistence in the human gut^34,35,37,79,80^. Estimates suggest that *Crassvirales* comprise > 86% of all gut phages, and *C. communis* alone comprises >40%^78^. In addition to their high abundance, *Crassvirales* are also persistent. *Crassvirales* can stably persist for long periods in the healthy human gut (>4 years), human gut post fecal microbiota transplants (>1 year), and cultures in the lab within a single bacterial host (>21 days)^21,34,72,76,79,81,82^. In lab culture, *Kehishuvirus primarius* (crAss001), the first cultured *Crassvirales*, is thought to exist in a carrier state infection,where fully formed phage particles are thought to be maintained within dividing cells without inducing cell lysis, thereby allowing its long-term persistence in culture in the absence of integrative lysogeny^76^. Additionally, DNA inversion of promoter sequences in the bacterial hosts of *Crassvirales* impacts the bacterial susceptibility to phage infection and dramatically reduces the ability of these phages to both persist and plaque^34,35,76^. Promoter inversion allows constant replenishment of resistant bacterial subpopulations via varied expression of cell surface structures implicated in the phage-bacterial binding interface. While the fact that only a small percentage of the bacteria population may be susceptible to phage receptor binding at any given time might be one possible explanation for the overall low efficiency of plaquing of the *Crassvirales,* we propose that non-integrative plasmid-like lysogeny might offer an alternative explanation. We hypothesize that plasmid lifestyles contribute to long-term persistence and low levels of host lysis as well as providing an escape to phase variable surface structures by eliminating the need to bind to them altogether.

The fact that *C. communis* does not plaque readily and in liquid culture does not seem to induce a growth defect in its host may suggest that it is in the process of losing its phage functions or that it maintains incredibly strict control over its lytic “on switch.” It is thought that PPs can lose their phage functions altogether once they obtain mobilization genes and an origin of transfer as an alternative method of horizontal transfer^83^. This could simply be due to genetic drift and pseudogenization of phage genes, or more likely due to the fact that PP genomes are larger than those of integrative phages and larger genome size leads to reduced efficiency of genome packaging into phage particles and a single large genetic acquisition event may inhibit packaging altogether^22,84^. *C. communis* does encode *mobC*, a mobilization gene, that when used with conjugative machinery encoded in the chromosome or another mobile element may enable cell-to-cell transfer of the PP genome without going through a lytic phase. This is one possible explanation for the lack of *C. communis* plaque formation observed. However, our results indicate that *C. communis* primarily replicates as a plasmid in the human gut, but phage-related functions are still transcribed. Our data supports a model in which *C. communis* primarily replicates as a plasmid, and uses a genetic switch to only lyse out of cells when its survival is threatened by its host population dwindling or when its bacterial host is under undue stress. We wonder if many *Crassvirales* are PPs and if this may explain the historical difficulty in culturing them and sampling their diversity.

Despite large culturing efforts, only eight *Crassvirales* of the >700 computationally identified *Crassvirales* genomes available in NCBI have been cultured, and of these, six form plaques and are therefore likely not representative of the *C. communis* lifecycle^34,35,37,79,80^. Two cultured *Crassvirales*, *Jahgtovirus secundus* (crAss002) and the unclassified “C4”, do not form plaques and thus studying them requires enrichment in liquid culture, which is resource-intensive and makes uncovering their biology difficult ^34,79^. While *J. secundus* persists in culture with its host, C4 biology is unknown and the studies of these two phages are limited, due to the laborious nature of culturing them^34,79^. Additionally, while these phages might be closer in lifestyle to the *C. communis* than the other cultured *Crassvirales,* no phages from the *C. communis* species or even the *Carjivirus* genus had been cultured prior to this study. With phage-targeted, plaque-free culturing approaches, we can start to bridge the gap between the computational discovery of novel viral genomes and the experimental characterization of their novel lifestyles. In this study, we demonstrate that targeted, plaque-free culturing can capture a PP, and we hypothesize that our method could also be used to culture other non-integrative lysogens and non-plaquing phages.

PPs are prolific agents of horizontal gene transfer, including implications in the spread of antibiotic resistance^39,85^. The spread of genetic content between bacteria within the gut may have dramatic effects on human health in terms of increased bacterial virulence, persistence, and antibiotic resistance^85,86^. Due to their higher levels of accessory genes and larger genomes there is more room for genome plasticity and genetic exchange without disrupting essential gene functions in PPs. PPs are also less drastically impacted by disruption of essential genes for one genetic program (phage or plasmid) because the alternate program may remain intact. In fact, transposable elements have been shown to frequently jump between PPs and plasmids^83^. These transposition events can catalyze the transition of one element type to another (plasmid to PP, PP to plasmid, PP to integrative phage, etc.)^83^. This raises a series of interesting questions that remain unexplored; how often are these transitions occurring? Is the frequency of these transitions driven by the fact that one lifestyle is more fit than another in the human gut? Is a non-integrative plasmid lysogen more fit than an integrative lysogen? While plasmids may be challenged by plasmid incompatibility, they can also be maintained at multiple copies per cell and their genes are more highly expressed than integrative lysogens that can only exist at one copy per cell and are often subjected to gene silencing, inactivation, and pseudogenization. However, integrative lysogens can more easily evade bacterial defense systems than extrachromosomal elements. More in depth study of these topics is required to understand the fitness advantages and disadvantages of each lifestyle type and the transitions between them.

Our study has several limitations. First, in the absence of plaque formation, obtaining a pure isolated stock of *C. communis* was not possible. Therefore, some of the growth dynamics we observe may be confounded by the replication of other phages in the cultures. Regardless, we do not observe detectable host lysis by OD600 and only observe minimal lysis by CFU on *P. vulgatus* and none on *P. dorei*, therefore lysis by additional phages in the culture is negligible. Second, while we did not observe plaque formation, it is possible that we simply were unable to identify the lytic trigger of *C. communis,* and that it is indeed capable of frequent and major lysis events. Given that excessive bacterial cell lysis can lead to disease states and inflammation, understanding the triggers of *C. communis* cell lysis may have large implications on human health^8,20^. However, the idea that *C. communis* predominantly replicates as a plasmid suggests that *C. communis* does not often impact human health through large shifts in microbial community composition via cell lysis in the absence of a lytic trigger. Further research in this area is required.

This study provides the first experimentally-backed insights into the biology and lifestyle of one of the most abundant and prevalent gut phages, for which we have very little understanding of its implications for human health. Many previous studies have attempted to correlate *C. communis* with different disease states, however in the absence of a confirmed bacterial host it was previously impossible to determine coordinated changes in phage and bacterial host abundance^87–91^. Generally, searches for phage-bacteria correlations rely on the assumption that replication of the phage is detrimental to its host. However, existence as a plasmid might allow *C. communis* to provide its bacterial host with fitness benefits, thereby shaping the microbial community composition via promoting cell growth rather than cell death. Further studies of *C. communis* are needed to gain an understanding of the complex relationship that it shares with both its microbial and human hosts.

## Acknowledgements

We thank Dylan Maghini for sharing Oxford Nanopore data, Jakob Wirbel for sharing his computational expertise, Michelle Hays for thoughtful discussions and experimental expertise, and Aravind Natarajan for thoughtful discussions on experimental design.

G.S. was supported by R35 GM131824. D.T.S. and A.S.H. were supported by the National Science Foundation Graduate Research Fellowship Program (DGE-1656518). D.T.S. was also supported by the Cell and Molecular Biology Training Grant (T32 GM007276). I.L. was supported in part by grants from NIAID (R44 AI172703, R44 AI162570) and a grant from the Bill & Melinda Gates Foundation. A.S.B. was supported by National Institutes of Health R01 AI148623, R01 AI143757, a Distinguished Investigator Award from the Paul Allen Foundation and the Stand Up 2 Cancer Foundation. Computing costs were supported, in part, by a NIH S10 Shared Instrumentation Grant S10 OD02014101.

## Author Contributions

D.T.S, A.S.B, and G.S. conceptualized the study. D.T.S. and A.S.H. performed all experiments; culturing of stool and isolation bacteria, qPCR. I.L. advised on ProxiMeta Hi-C sequencing and subsequent analysis. D.T.S. performed all computational analyses. D.T.S., G.S., and A.S.B wrote this manuscript with input from all authors.

## Methods

### Identification of other genes with plasmid origin in the *C. communis* genome (TA system and *mobC*)

To identify plasmid-like genes in the *C. communis* genome, we created a custom BLAST database of protein genes by downloading all of the protein sequences from plasmids in NCBI. Protein sequences from the *C. communis* reference genome (NCBI NC_067194) were compared against the custom database by running the command line version of blastp^92^. Hits were filtered for at least 50% query coverage and 25% identity. Hits that met this threshold were then aligned to their top blastp hit and a few additional related sequences using Geneious (Geneious, Muscle, and Clustal Omega alignment tools) (Geneious Prime 2023.2.1). If query sequences were less than 60% of the length of their closest hit, or vice versa, they were not considered.

### Identification of *repL* genes in other *Crassvirales*

To identify potential *repL* genes in other *Crassvirales* genomes, we used the RepL Pfam profile (PF01719) and searched all downloaded from NCBI using hmmsearch in HMMER 3.4^93^. Genomes from which identified proteins originated were blasted against the NCBI non-redundant protein sequences (nr) database to determine the similarity to other *Crassvirales*.

### Identifying *C. communis* containing stool

Stool previously sequenced via the shotgun illumina platform with paired 150bp reads was analyzed, and is available on SRA with the bioproject number PRJNA707487^62^. Kraken2^94^ was used for read classification using a custom database as previously described in^62^ and samples were first filtered by presence of the *Carjivirus* genus (>1% of reads classified). In samples with a high percentage of the *Carjivirus* genus, Bowtie2 2.5.4^95,96^ was used to align reads to the *C. communis* reference genome (NCBI NC_067194).

### Meta-Hi-C sequencing of *C. communis* containing stool and assembly of the *C. communis* genome

Meta-Hi-C sequencing was performed using the ProxiMeta kit provided by Phase Genomics exactly as directed in the standard protocol. 150 bp paired end reads were generated (Novaseq Illumina) as suggested by Phase Genomics’ protocols. The Phase Genomics computational ProxiMeta pipeline was used for assembly of the *C. communis* genome.

### Identification of bacterial host in meta-Hi-C sequencing and generation of interaction maps

The Phase Genomics computational ProxiMeta pipeline was used for counting chimeric reads between *C. communis* and bacterial assemblies from the sample. Hi-C interaction maps were generated with distiller default settings ^97^. Plots were visualized using python cooltools^98^ and matplotlib^99^.

### Computational host prediction via BACON homology and CRISPR spacer analysis

BACON domain homology searching was performed by BLASTing the translated sequence of the BACON domain containing protein sequence from the *C. communis* reference genome with blastp^92^. The tree file was downloaded from the BLAST results and the tree was built using ggtree in Rstudio ^100^. CRISPR spacer analysis was performed using Phisdetector ^64^.

### Oxford Nanopore sequencing of *C. communis* containing stool

Oxford Nanopore libraries were prepared with Oxford Nanopore ligation sequencing kit SQK-LSK109 and sequenced on one FLO-MIN106 flow cell. Reads were assembled with Lathe v1.0^101^; briefly, reads were basecalled then assembled into contigs with Canu^102^, and contigs were polished with short reads^103^.

### Analysis of Oxford Nanopore sequencing for *C. communis* integration events

Oxford Nanopore sequencing reads and assembled contigs were mapped to the *C. communis* reference genome using minimap2 2.26-r1175^104^. Regions of clipped reads were extracted using samtools v1.19^105^ and aligned back to the *C. communis* assembled contig using minimap2 2.26-r1175^104^. Clipped regions that did not align back to the *C. communis* assembly were BLASTed against the standard nucleotide database (nucleotide collection (nr/nt)) using blastn^92^.

### Origin of replication prediction

Origins of replication were predicted with OriFinder-2022^71^. Oxford Nanopore data were mapped to the *C. communis* reference genome via minimap2 2.26-r1175^104^. GC skew plot was generated with SkewIT^106^.

### Analysis of publicly available metatranscriptomics data

Paired metagenomics and metatranscriptomics data were downloaded from SRA for the bioproject PRJNA354235^107^. Reads were aligned to the *C. communis* reference genome using Bowtie2^95^. Samples with at least 300 reads (∼1x coverage) aligning to *C. communis* were used for downstream analyses (n = 111). Bedtools v2.27.1 coverage was used to calculate the coverage for each gene in the genome^108^. Gene length was determined in kilobases. Per million scaling factor was determined by dividing the number of reads mapping to *C. communis* by 1,000,000. Next, RPM was calculated; RPM = read counts for a gene / per million scaling factor. Finally, RPKM was calculated; RPKM = RPM / gene length.

### Comparison of *Crassvirales* genomes and generation of synteny plots

To compare the similarity of our *C. communis* assembly to the *C. communis* reference genome we used the Geneious mapper with default parameters (Geneious Prime 2023.2.1). To compare the genomes of all of the cultured species of *Crassvirales* we performed whole genome alignment using Clustal Omega with default settings^109^. From the clustal omega output we constructed a phylogenetic tree using the Geneious tree builder (Geneious Prime 2023.2.1) with default parameters. We visualized similarity in gene organization and produced gene plots of the genomes using AnnoView^110^. Synteny plots were generated using EasyFig 3.0^111^. NCBI reference numbers of genomes are as follows; *C. communis* NC_067194, *R. jaberico* OQ198719, *K. frurule* OQ198718, *K. tikkala* OQ198717, DAC15 NC_055832, crAss001 NC_049977, crAss002 MN917146, 14:2 KC821624. C4 genome is not available on NCBI and was downloaded from the supplement of the paper^79^.

### Stool-based culture

1 mg of stool frozen in no preservatives at −80°C was resuspended in 1 mL of Brain heart infusion (BHI) liquid media. The stool suspension was vigorously vortexed until homogenized. Homogenized stool resuspension is diluted 1:100 into anaerobic tryptic soy broth (TSB) containing vancomycin (7.5 μg/ml) and kanamycin (100 μg/mL). The stool was diluted into 50 mL of TSB and cultured at 37°C anaerobically for 44 hours. Anaerobic culturing was performed in an anaerobic chamber (Bactron).

### Quantitative PCR assays

qPCR reactions were set up in technical duplicate 10 µL reactions in 384-well plates. Standard curves were constructed using plasmids with the target sequences cloned into them and diluted tenfold. Plasmids were ordered through IDT via synthesis of the target sequence and cloning it into the IDTsmart backbone. Reactions were set up according to standard protocols for the Applied Biosystems Power SYBR Green PCR Master Mix. *C. communis* targeting primers were previously published^112^, crAss056_F (CAGAAGTACAAACTCCTAAAAAACGTAGAG) and crAss056_R (GATGACCAATAAACAAGCCATTAGC). *P. vulgatus* targeting primer sequences were Pv_gmk_F (GGAAAAGAACGGCATGGTGT) and Pv_gmk_R (ATCCGCCTACCACATCTACG), and were designed to target the guanylate kinase (*gmk*) gene.

### Culturing phage filtrate on predicted bacterial hosts

After 44 hours of stool-based culture the viral fraction was harvested by pelleting the bacteria and filtering the supernatant through a 0.2 µm filter. The bacterial hosts were grown in anaerobic overnight cultures of Brain Heart Infusion Supplemented with Hemin and Cysteine (BHIS) at 37°C. The overnight cultures were then diluted 1:50 into fresh anaerobic BHIS and grown for roughly four hours until they reached an OD600 reading of ∼0.1-0.3. Phage filtrate was added to bacterial culture with an MOI of ∼0.1 based on qPCR quantification of viral copies. Samples were split into three replicates. As controls, bacterial cultures were grown without adding phage and phage filtrate was added to fresh BHIS media. Cultures were anaerobically incubated at 37°C for 44 hours. The *P. vulgatus* strain was obtained from ATCC (*Phocaeicola vulgatus* ATCC 8482^TM^). The *P. dorei* strain was isolated from stool, whole genome sequenced with paired 150bp reads (Novaseq Illumina).

### Analysis of publicly available single-cell microbiome sequencing data

Publicly available single-cell microbiome sequencing data were downloaded from SRA bioproject PRJNA803937^63^ and aligned to the *C. communis* reference genome (NC_067194) and the *P. vulgatus* genome (*Phocaeicola vulgatus* ATCC 8482TM) using Bowtie2 2.5.4^95^. The same 14 samples identified in the original publication with at least 5% of reads mapping to *C. communis* infecting *P. vulgatus* were identified. Coverage of each genome (*C. communis* and *P. vulgatus*) was calculated using bedtools^108^ genomecov by base. The coverage of each base in the genome was summed and divided by the genome length to calculate the average coverage of the genome. The ratio of *C. communis* to *P. vulgatus* coverage was taken by dividing the *C. communis* coverage from each sample by the *P. vulgatus* coverage.

## Supplementary Figure Legends

**Supplementary Fig 1. *repL* genes are found in other *Crassvirales***

Protein alignments of all RepL proteins found in *Crassvirales*. A) phylogeny based on protein alignments. 23 protein sequences are found across 19 *Crassvirales* genomes. Black *s denote 9 proteins across 8 genomes of *Crassvirales* species that share >70% nucleotide identity across >88% of their genomes, but share no similarity to *C. communis.* Teal *s denote 4 proteins across 4 divergent genomes of *Crassvirales* (divergent both from on another and *C. communis*). Gray *s denote 5 proteins across 4 *Crassvirales* genomes that share >96% nucleotide identity across >96% of the *C. communis* reference genome. Blue *s denote 2 proteins in one *Crassvirales* genome belonging to the *Carjivirus* genus. Bright green *s denote 3 proteins across 2 *Crassvirales* genomes belononging to *Intestiviridae.* B) protein alignments of RepL sequences that correspond to the tree shown in part A.

**Supplementary Fig 2. ProxiMeta Hi-C sequencing contact maps between *C. communis* and bacterial MAGs**

A/B) ProxiMeta Hi-C maps of *P. vulgatus* and *C. communis* links at two different heat scales, heat map color represents number of ProxiMeta Hi-C links. C/D) ProxiMeta Hi-C maps of P. merdae and *C. communis*. E/F) ProxiMeta Hi-C maps of *B. stercoris* and *C. communis* G/H) ProxiMeta Hi-C maps of an unclassified *Prevotella* species and *C. communis*.

**Supplementary Fig 3. ProxiMeta Hi-C data suggests that *C. communis* does not integrate into *P. vulgatus***

A) Number of ProxiMeta Hi-C links (y-axis) across 10kb bins. Bins 1-10 are *C. communis*. Bins >10 are *P. vulgatus*. Mann-Whitney U Test: mean for bins 10 and lower (*C. communis*): 563.9, mean for bins 11 and larger (*P. vulgatus*): 0.859, P-value: 8.766e-09, means are statistically significantly different. B) zoom in on part A, only showing *P. vulgatus* bins. z-test to determine outliers based on their deviation from the mean did not determine any outliers.

**Supplementary Fig 4. Expression of *C. communis* genes in metatranscriptomics**

A) Count of samples with RPKM >0 for each gene in *C. communis*, plasmid related genes in green, phage related genes in black. B) Average RPKM for each gene in *C. communis* across all samples (n = 111), plasmid related genes in green, phage related genes in black.

**Supplementary Fig 5. *C. communis* has a wide host range and does not impact OD600**

A) OD600 of *P. vulgatus* culture alone (purple) or with phage added (green). OD600 of media only (black) and media with phage added (gray). B) OD600 of *P. dorei* culture alone (purple) or with phage added (green). OD600 of media only (black) and media with phage added (gray). C) OD600 of *B. stercoris* culture alone (purple) or with phage added (green). OD600 of media only (black) or media with phage added (gray). D) OD600 of *P. merdae* culture alone (purple) or with phage added (green). OD600 of media only (black) or media with phage added (gray). E) Copies of *C. communis* per mL of culture were measured via qPCR in *P. merdae* culture (purple), in *B. stercoris* culture (green), and in media only (gray). *C. communis* was undetectable in cultures where phage was not added to *P. merdae*, *B. stercoris*, and media.

**Supplementary Fig 6. *C. communis* qPCR standard curves**

Samples were run across multiple qPCR plates, and a separate standard curve was run on each plate. We plotted the *C. communis* standard curve for each plate A-D) standard curves of *C. communis* cultured with isolated bacteria time courses. E-G) standard curves of *C. communis* standards for stool-based culture time course. H) all *C. communis* standards plotted together, dotted lines represent mean standard curve and +/- 3 Cq (shifted ∼1 log in each direction).

**Supplementary Fig 7. *P. vulgatus* qPCR standard curves**

A-B) standard curves of *P. vulgatus* standards for stool-based culture time course. C) all *P. vulgatus* standards plotted together, dotted lines represent mean standard curve and +/- 3 Cq (shifted ∼1 log in each direction).

**Supplementary Fig 8. *C. communis* qPCR replicates**

qPCR technical replicates where the qPCR primers target *C. communis* for A) media with phage filtrate added (-phage filtrate, copies of *C. communis* are undetected) B) *P. merdae* + phage filtrate (-phage filtrate, copies of *C. communis* are undetected) C) *B. stercoris* + phage filtrate (-phage filtrate, copies of *C. communis* are undetected) D) *P. dorei* + phage filtrate (-phage filtrate, copies of *C. communis* are undetected) E) *P. vulgatus* + phage filtrate (-phage filtrate, copies of *C. communis* are undetected) F) stool-based culture (-phage filtrate, copies of *C. communis* are undetected)

**Supplementary Fig 9. *P. vulgatus* qPCR replicates**

qPCR technical replicates for stool-based culture, where the qPCR primers target *P. vulgatus*.

---

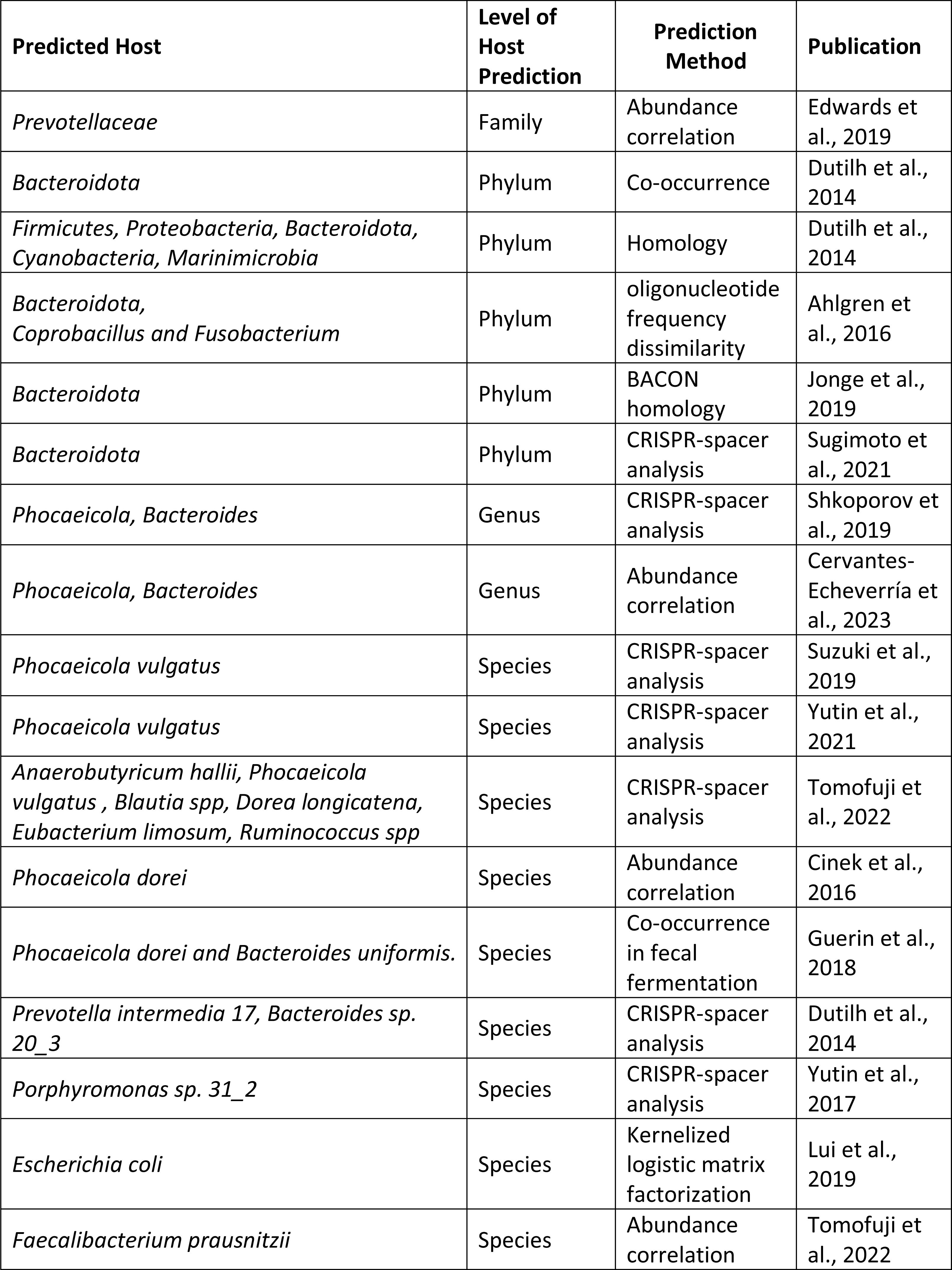
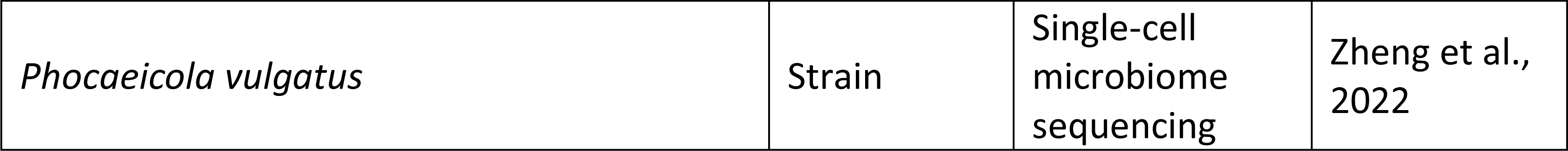

| Predicted Host | Level of Host Prediction | Prediction Method | Publication |
| --- | --- | --- | --- |
| *Prevotellaceae* | Family | Abundance correlation | Edwards et al., 2019 |
| *Bacteroidota* | Phylum | Co-occurrence | Dutilh et al., 2014 |
| *Firmicutes, Proteobacteria, Bacteroidota, Cyanobacteria, Marinimicrobia* | Phylum | Homology | Dutilh et al., 2014 |
| *Bacteroidota, Coprobacillus and Fusobacterium* | Phylum | oligonucleotide frequency dissimilarity | Ahlgren et al., 2016 |
| *Bacteroidota* | Phylum | BACON homology | Jonge et al., 2019 |
| *Bacteroidota* | Phylum | CRISPR-spacer analysis | Sugimoto et al., 2021 |
| *Phocaeicola, Bacteroides* | Genus | CRISPR-spacer analysis | Shkoporov et al., 2019 |
| *Phocaeicola, Bacteroides* | Genus | Abundance correlation | Cervantes-Echeverría et al., 2023 |
| *Phocaeicola vulgatus* | Species | CRISPR-spacer analysis | Suzuki et al., 2019 |
| *Phocaeicola vulgatus* | Species | CRISPR-spacer analysis | Yutin et al., 2021 |
| *Anaerobutyricum hallii, Phocaeicola vulgatus, Blautia spp, Dorea longicatena, Eubacterium limosum, Ruminococcus spp* | Species | CRISPR-spacer analysis | Tomofuji et al., 2022 |
| *Phocaeicola dorei* | Species | Abundance correlation | Cinek et al., 2016 |
| *Phocaeicola dorei and Bacteroides uniformis.* | Species | Co-occurrence in fecal fermentation | Guerin et al., 2018 |
| *Prevotella intermedia 17, Bacteroides sp. 20_3* | Species | CRISPR-spacer analysis | Dutilh et al., 2014 |
| *Porphyromonas sp. 31_2* | Species | CRISPR-spacer analysis | Yutin et al., 2017 |
| *Escherichia coli* | Species | Kernelized logistic matrix factorization | Lui et al., 2019 |
| *Faecalibacterium prausnitzii* | Species | Abundance correlation | Tomofuji et al., 2022 |
| *Phocaeicola vulgatus* | Strain | Single-cell<br>microbiome<br>sequencing | Zheng et al.,<br>2022 |

