## Supplementary figures and images for "Analysis and culturing of the prototypic crAssphage reveals a phage-plasmid lifestyle"

### Supplementary Figure 1

#### Supplementary figure 1

A

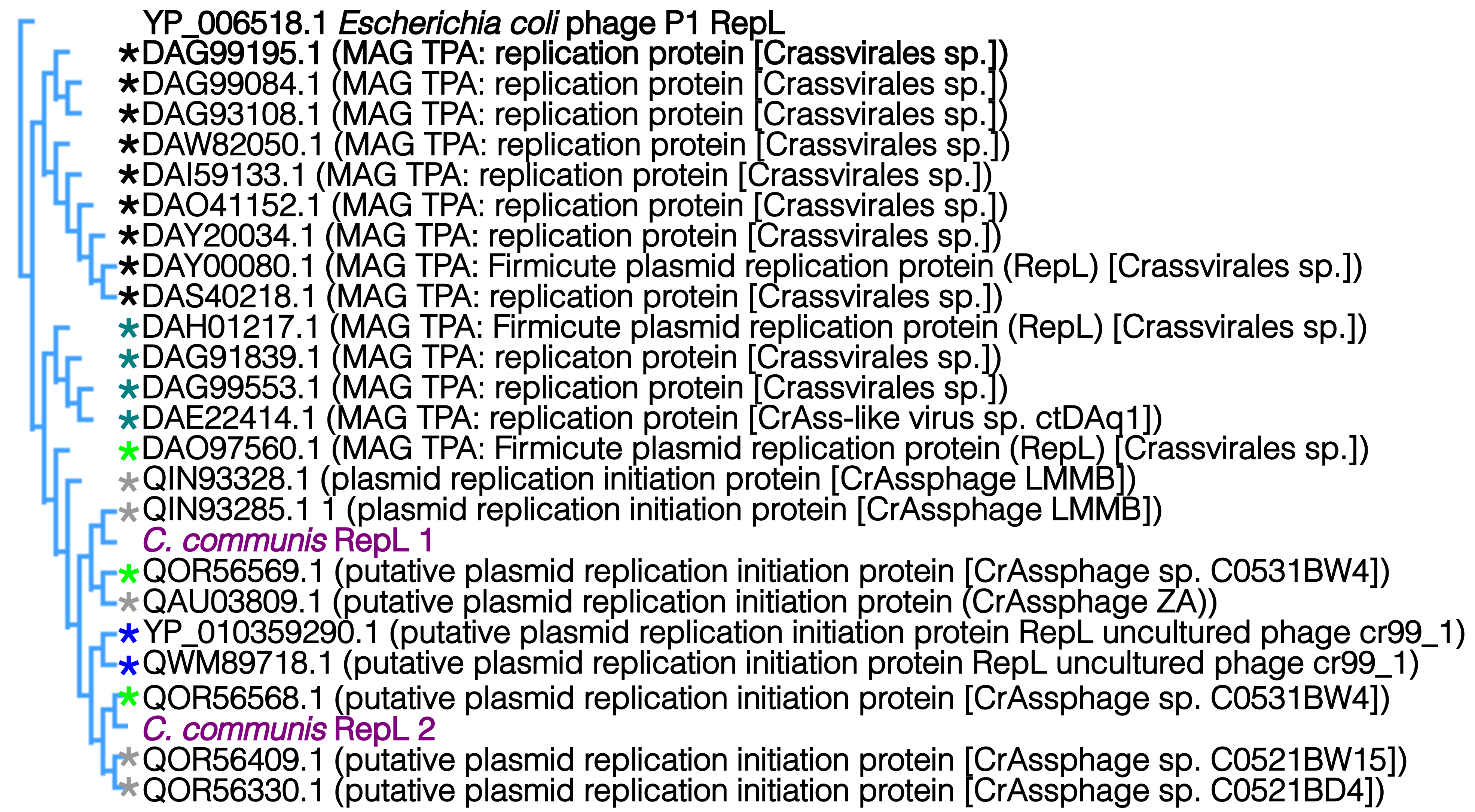


B

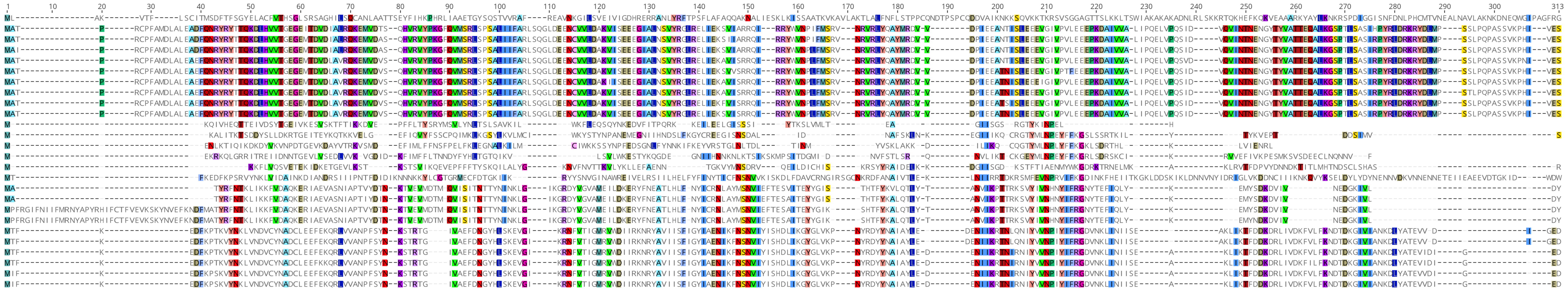

### Supplementary Figure 2

#### Supplementary figure 2

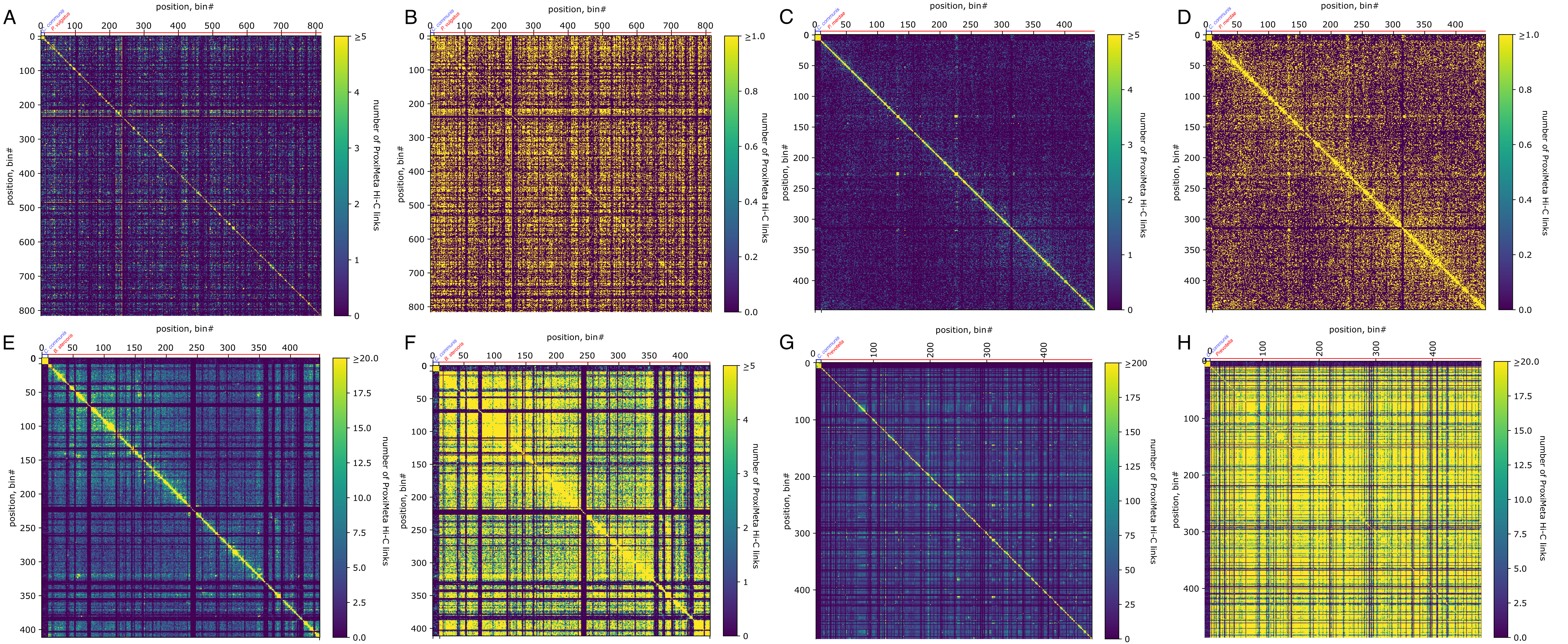

### Supplementary Figure 3

#### Supplementary figure 3

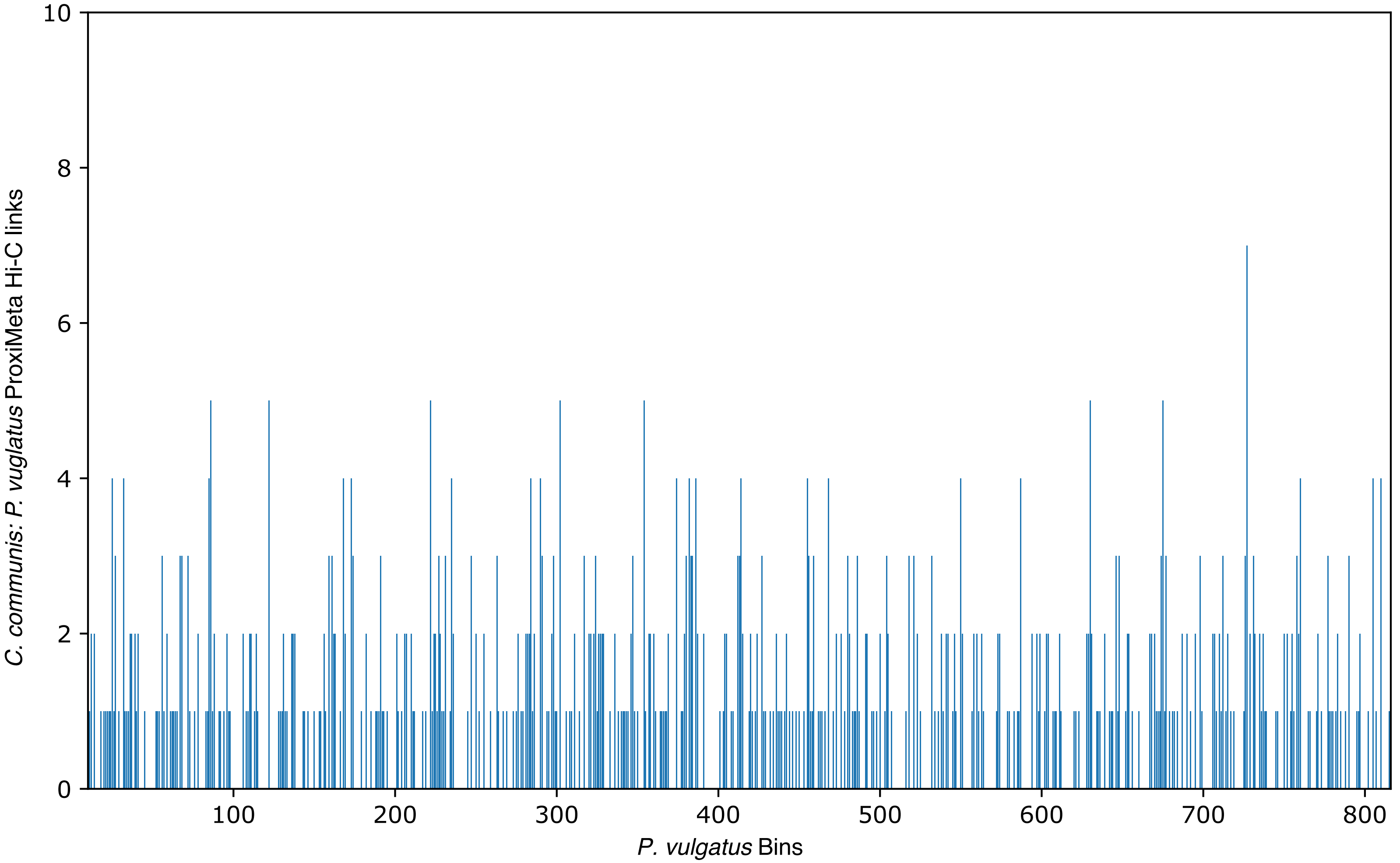

### Supplementary Figure 4

#### Supplementary figure 4

A

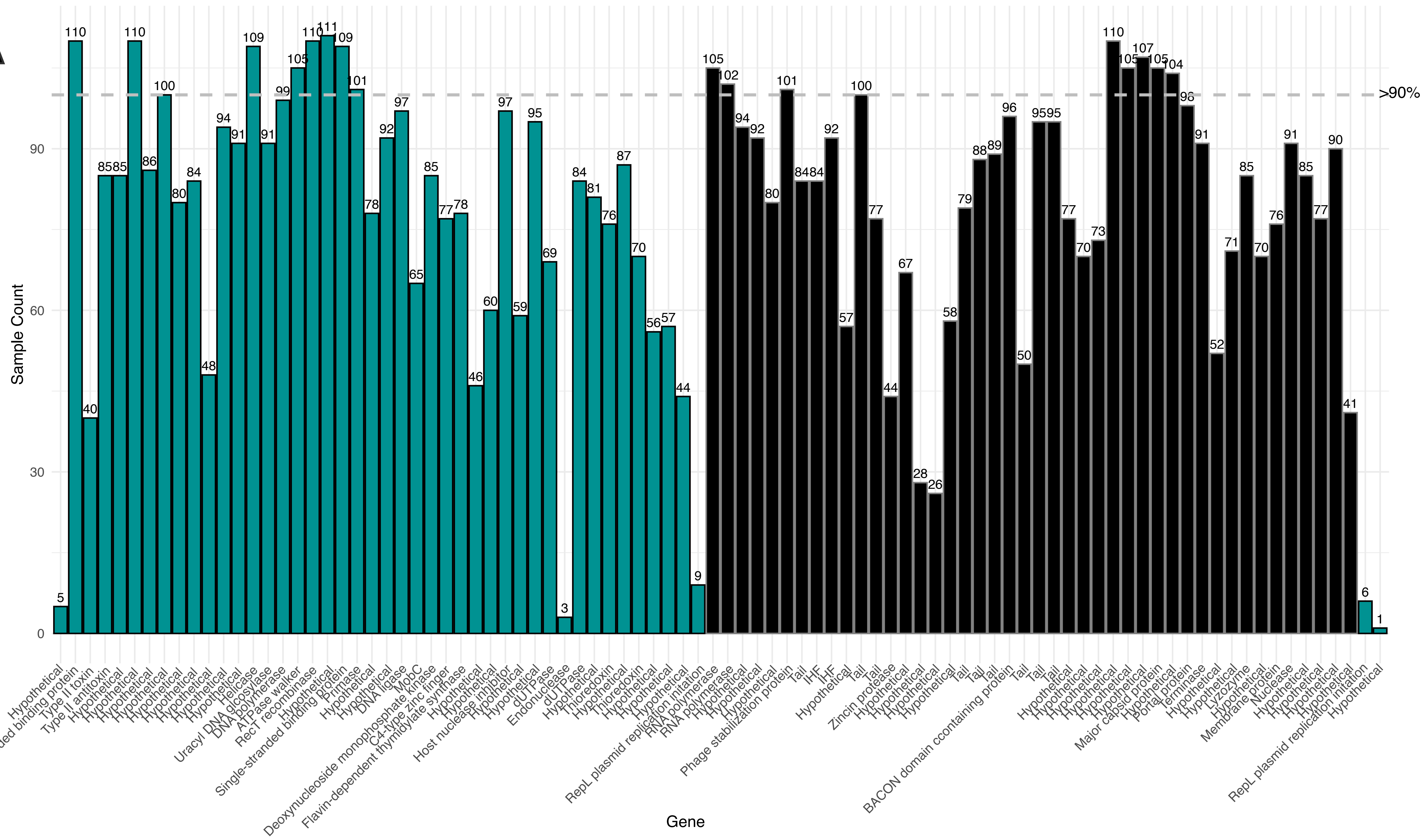


B

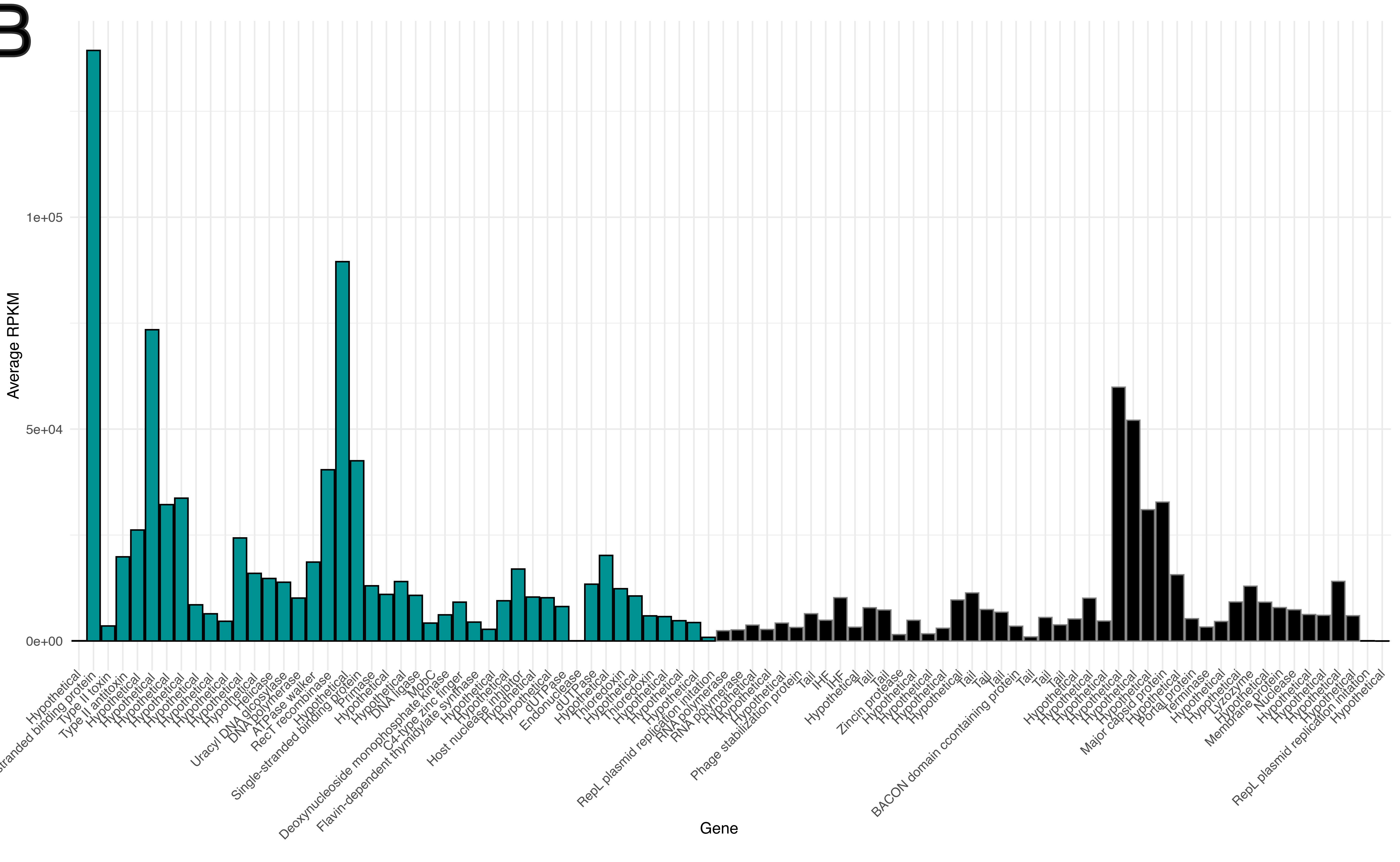

### Supplementary Figure 5

#### Supplementary figure 5

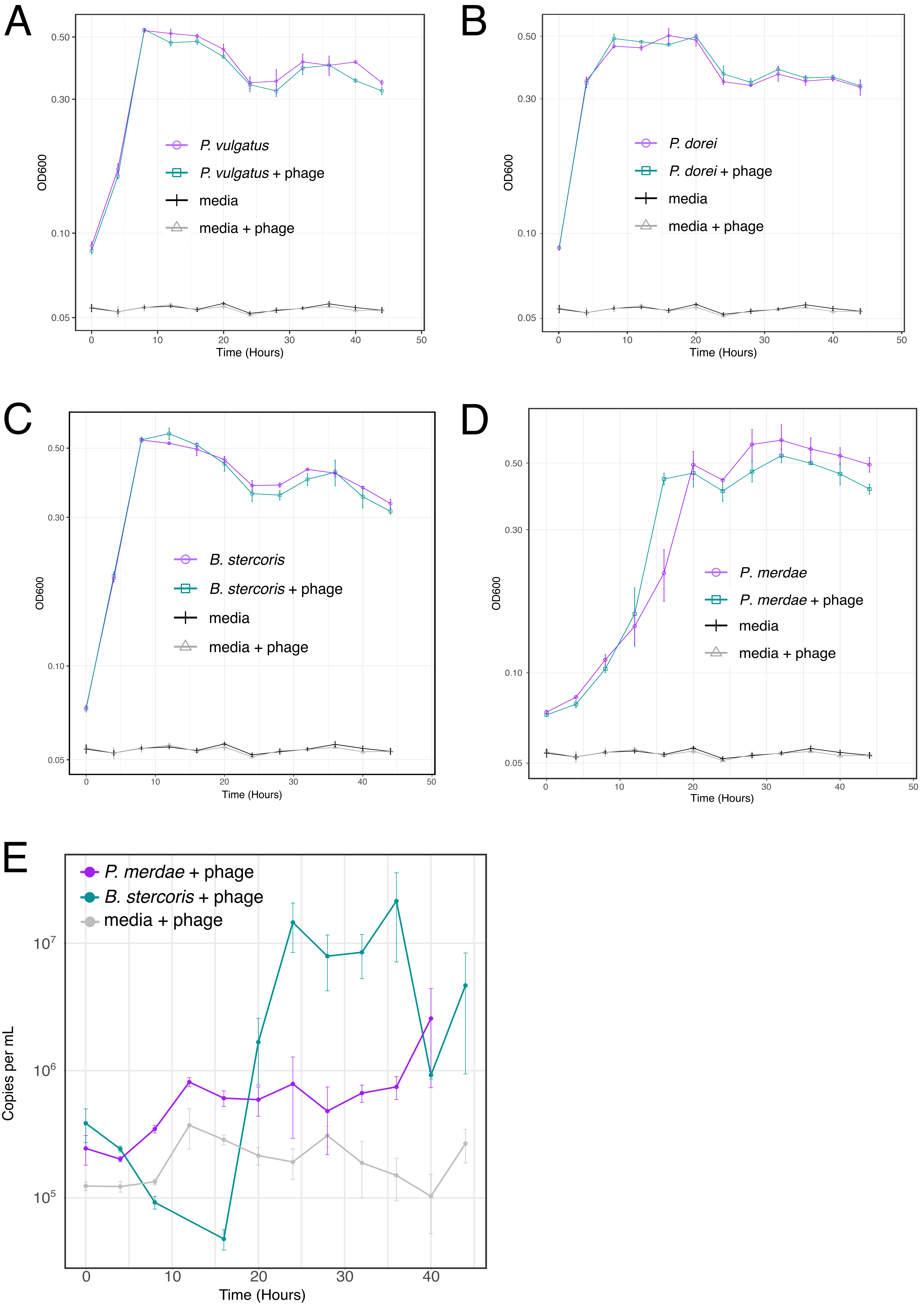

### Supplementary Figure 6

#### Supplementary figure 6

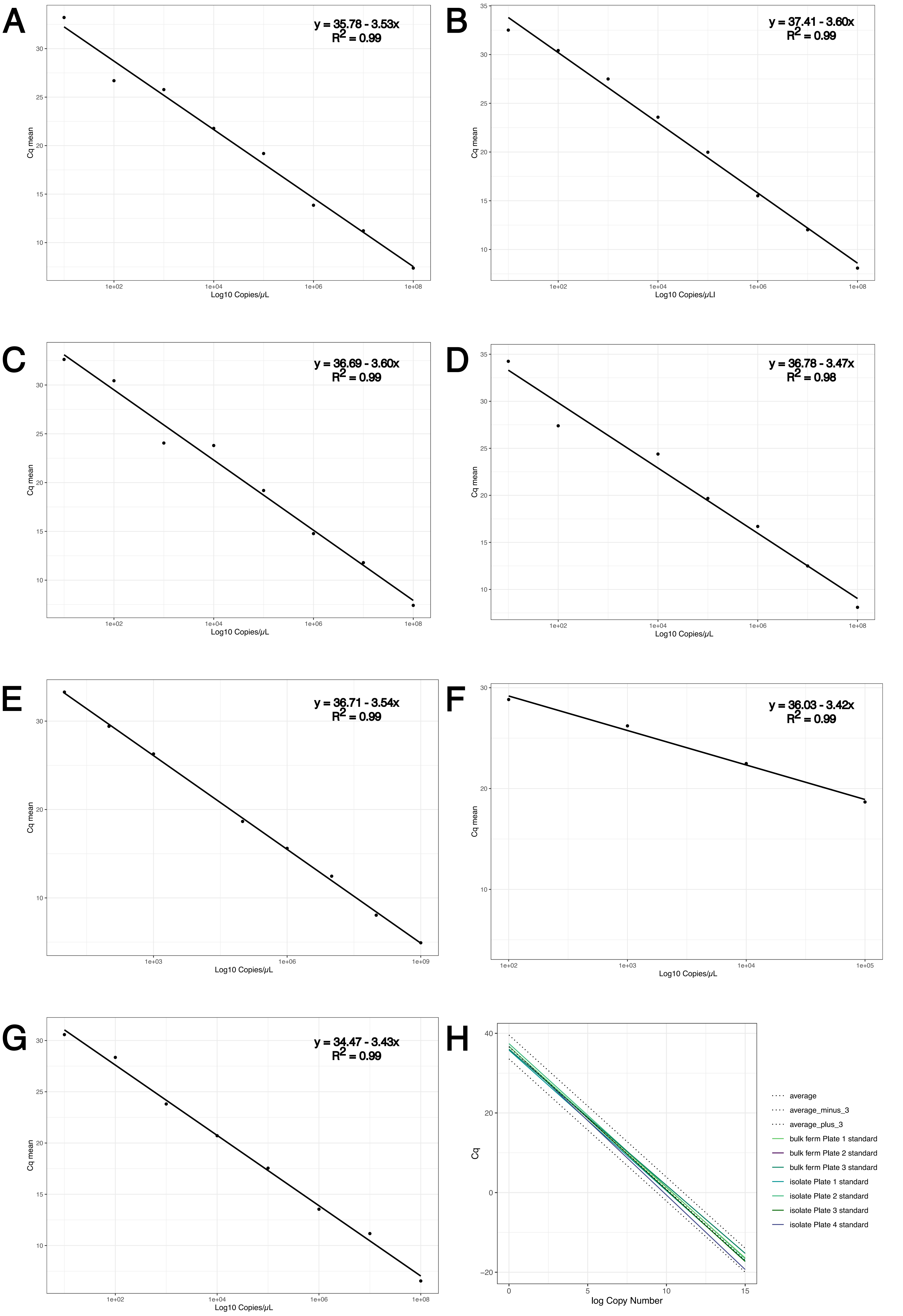

### Supplementary Figure 7

#### Supplementary figure 7

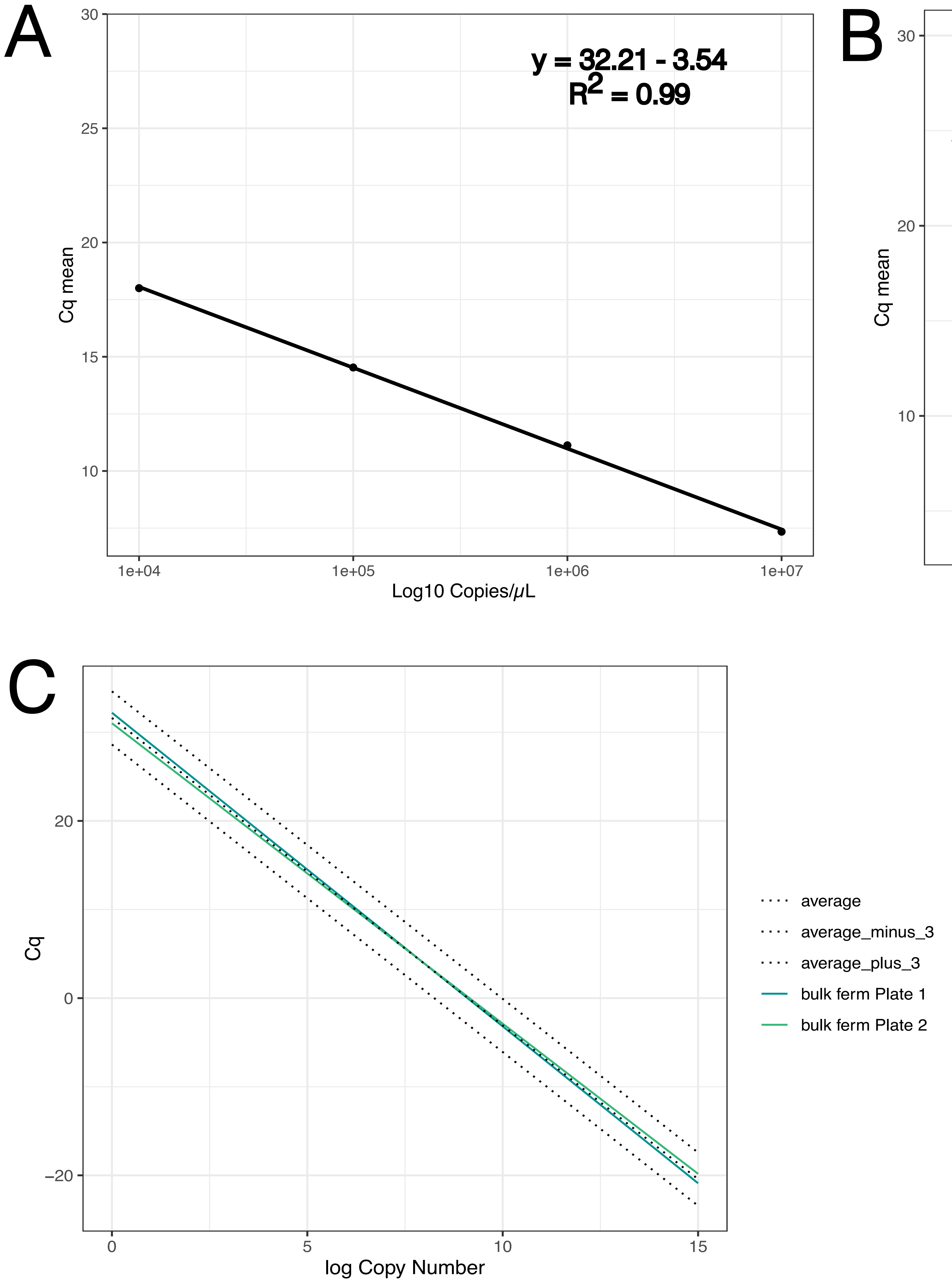

### Supplementary Figure 8

#### Supplementary figure 8

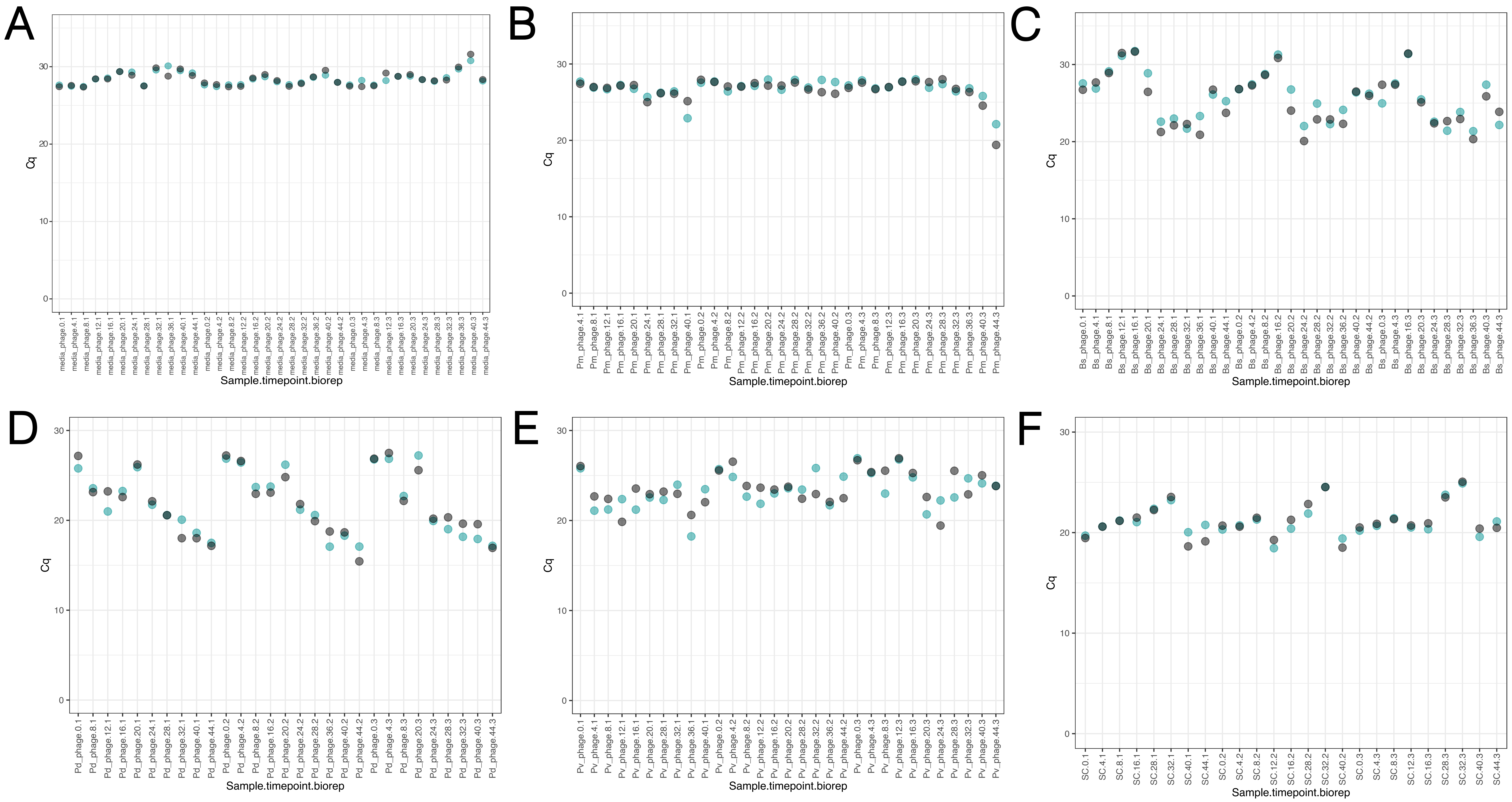

### Supplementary Figure 9

#### Supplementary figure 9

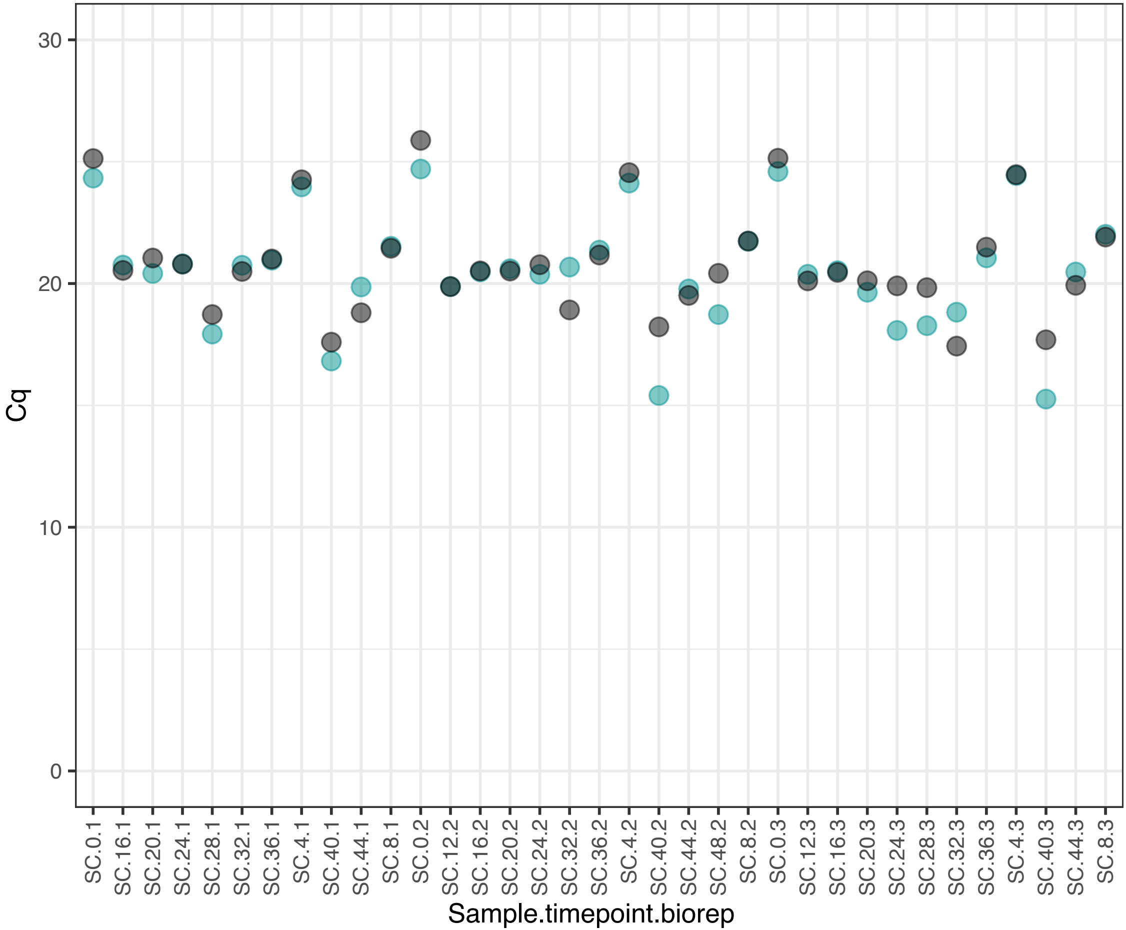
